# Role of Complementarity-Determining Regions 1 and 3 in Pathologic Amyloid Formation by Human Immunoglobulin κ1 Light Chains

**DOI:** 10.1101/2023.02.01.526662

**Authors:** Elena S. Klimtchuk, Daniele Peterle, Esther A. Bullitt, Lawreen H. Connors, John R. Engen, Olga Gursky

## Abstract

Immunoglobulin light chain (LC) amyloidosis is a life-threatening disease whose understanding and treatment is complicated by vast numbers of patient-specific mutations. To address molecular origins of the disease, we explored 14 patient-derived and engineered proteins related to κ1-family germline genes IGKVLD-33*01 and IGKVLD-39*01. Hydrogen-deuterium exchange mass spectrometry analysis of local conformational dynamics in full-length recombinant LCs and their fragments was integrated with studies of thermal stability, proteolytic susceptibility, amyloid formation, and amyloidogenic sequence propensities using spectroscopic, electron microscopic and bioinformatics tools. The results were mapped on the atomic structures of native and fibrillary proteins. Proteins from two κ1 subfamilies showed unexpected differences. Compared to their germline counterparts, amyloid LC related to IGKVLD-33*01 was less stable and formed amyloid faster, whereas amyloid LC related to IGKVLD-39*01 had similar stability and formed amyloid slower. These and other differences suggest different major factors influencing amyloid formation. In 33*01-related amyloid LC, these factors involved mutation-induced destabilization of the native structure and probable stabilization of amyloid. The atypical behaviour of 39*01-related amyloid LC tracked back to increased dynamics/exposure of amyloidogenic segments in βC’_V_ and βE_V_ that could initiate aggregation, combined with decreased dynamics/exposure near the Cys23-Cys88 disulfide whose rearrangement is rate-limiting to amyloidogenesis. The results suggest distinct amyloidogenic pathways for closely related LCs and point to the antigen-binding, complementarity-determining regions CDR1 and CDR3, which are linked via the conserved internal disulfide, as key factors in amyloid formation by various LCs.

## Introduction

Amyloid light chain (AL) amyloidosis is the most common human systemic amyloid disease worldwide [1, 2]. This aggressive lethal disease results from misfolding and aggregation of immunoglobulin (Ig) light chains (LCs) overproduced in the course of monoclonal β-cell dyscrasia. Diverse manifestations of AL amyloidosis involve deposition of fibrillary LCs and their fragments [3, 4] in vital organs such as kidney and heart, causing organ damage and failure (reviewed in [2, 5, 6]). These and other pathologies arise from excess circulating monoclonal LCs with particular physicochemical properties conferred by amino acid substitutions in variable domains (V_L_). Identifying these properties and their links to proteotoxicity has been particularly challenging due to the vast numbers of V_L_ mutations that lead to natural diversity of LC sequences, their misfolding pathways and amyloid polymorphs ([2, 7-9] and references therein).

Of all LC λ and κ sub-families, λ6 and κ1 are predominant in AL amyloidosis [9-11]. Limited structural information is available on amyloids comprised of κ LCs [12], while λ LCs have been explored in greater detail. Early misfolding and aggregation intermediates have been characterized biophysically for λ3 V_L_ [8]. Importantly, *ex vivo* amyloid fibril structures have been determined by cryo-electron microscopy (cryo-EM) to near-atomic resolution for λ1, λ3 and λ6 LCs, revealing different amyloid polymorphs for individual LCs [9, 13-15]. Despite these differences, all AL-related fibril structures available to date share several common features: i) the amyloid core is derived from V_L_ segments; ii) the flattened molecular shape in amyloid fibrils bears no similarity to the native fold; and iii) fibril formation involves an ∼180° rotation around the conserved disulfide that links parallel β-strands B_V_ and F_V_ (βB_V_ and βF_V_) in native V_L_; these segments become antiparallel in the fibril core. Such a polypeptide chain rotation, which must involve transient unfolding of the native structure around the internal disulfide, was proposed to be rate-limiting in amyloid formation [14, 15].

Native structures of LCs are comprised of two β-sandwich Ig domains, variable and constant (C_L_), joined via the J-region (Fig. 1); variable-joined region is termed V_L_ hereafter. Three complementarity-determining regions (CDRs) in V_L_ form the antigen binding loops. In circulation, LCs readily self-assemble into noncovalent dimers [16]. Dissociation of these dimers, followed by the proteolytic cleavage downstream of the J-region to release the misfolding-prone V_L_ fragments was proposed to generate the protein precursors of amyloid. However, the causal role of such cleavage in disease development is unclear, as full-length LCs are also found in amyloid deposits [3, 17]. Moreover, mass spectrometry studies of AL deposits suggest that LC domain separation can follow the misfolding rather than precede it [4, 18], and the exact sequence of steps leading to formation of AL deposits remains to be determined.

**Figure 1.**
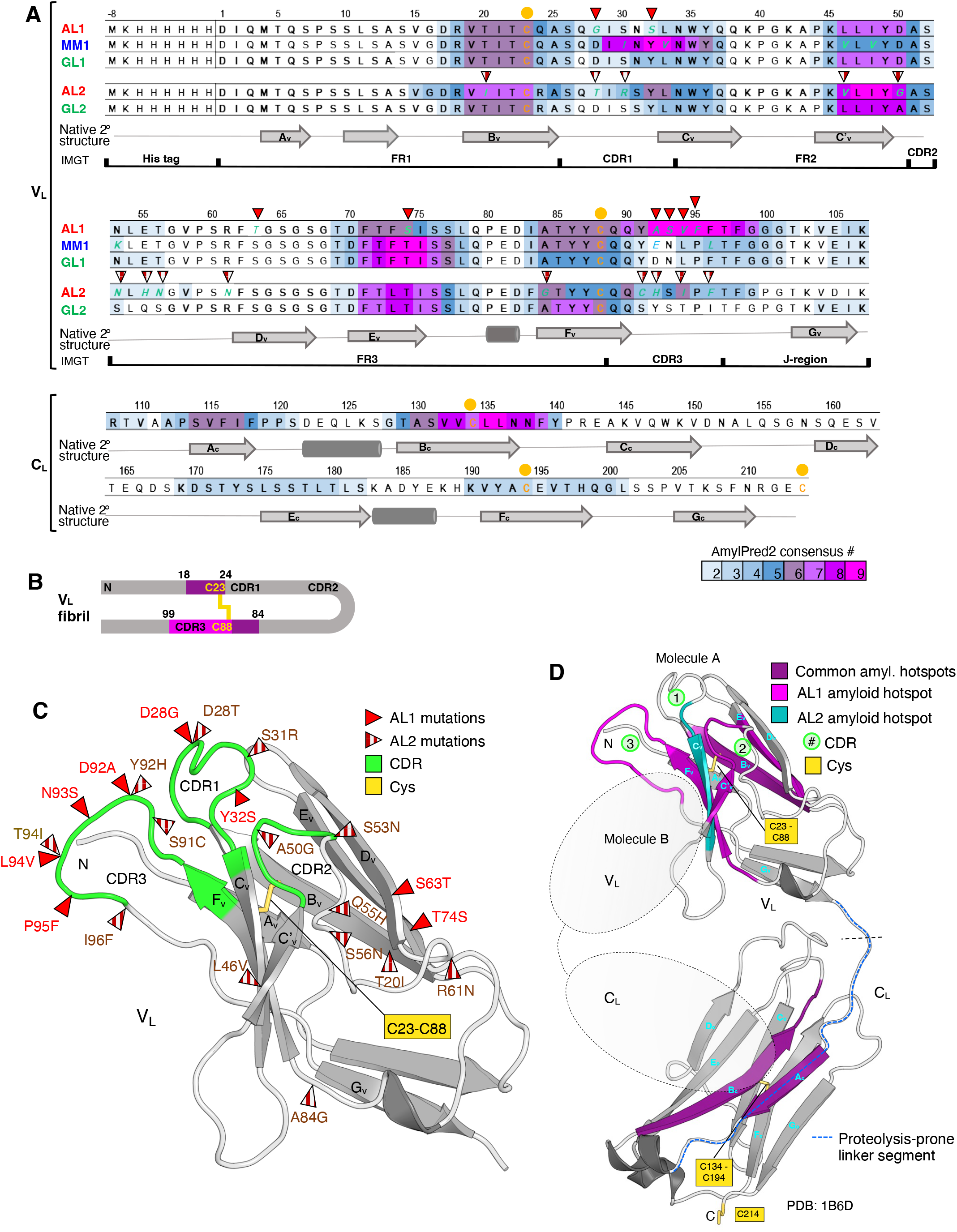
Structures and amyloid-forming propensities of κ1 LCs explored in the current study. **(A)** Amino acid sequences of LCs related to germline genes *IGKVLD-33*01* (AL1, MM1 and GL1) and *IGKVLD-39*01* (AL2 and GL2). Residues in AL and MM that differ from their GL counterparts are in green italics. Of the five cysteines (orange dots), C23-C88 and C134-C194 form internal disulfides, while C214 is either free or disulfide-linked to another stand-alone LC. The amyloidogenic sequence propensity predicted using AmylPred2 [28] is color-coded according to the consensus number, which increases with increased amyloidogenic sequence propensity. Framework and complementarity-determining regions (FRs and CDRs) and the J-region are indicated. Linear diagram shows secondary structure in the native protein based on the X-ray crystal structure of κ1 LC (PDB ID 1B6D) [36]; β-strands (A_V_ – G_V_ in V_L_ and A_C_ – G_C_ in C_L_) are marked. **(B)**Schematics illustrating the polypeptide chain orientation in V_L_ of the amyloid core. The internal disulfide C23-C88 and CDRs are indicated. Amyloid hotspots located in or near CDR1 and CDR3 in AL1 LC of the current study are shown; hotspots were predicted either in AL1 LC only (pink) or both in AL1 and GL1 LCs (violet). **(C, D)** Representative 3D structure of κ1 full-length LC and its V_L_ domain alone. β-strands and cysteines are marked. Locations of AL substitutions in V_L_ are shown (full red triangles for AL1, white and red triangles for AL2). CDRs (green) and the proteolysis-prone linker segment (blue dotted line) are shown in LC. Amyloid hotspots with AmylPred2 values equal to or greater than 5 that are in common between AL1 and AL2 are highlighted in purple, while those exclusively present in AL1 or AL2 are marked in bright purple or teal, respectively. The approximate location of the second LC molecule (Molecule B) within the LC dimer is schematically shown with two gray ovals.

Like other amyloidogenic globular proteins, AL LCs often show decreased thermodynamic and kinetic stability as compared to their non-amyloidogenic counterparts, yet decreased stability alone is neither necessary nor sufficient for amyloid deposition [17, 19, 20]. Other proposed drivers of AL amyloidosis can include: i) destabilization of the LC dimer due to weakened V_L_-V_L_ and C_L_-C_L_ interactions [21, 22]; ii) destabilization of the full-length LC due to weakened V_L_-C_L_ interactions that are regulated, in part, by the inter-domain linker downstream of the J-region [23]; and iii) specific V_L_ segments, such as the cleavage-prone V_L_-C_L_ linker [18, 23] or “sticky” segments with high amyloid-forming sequence propensity, aka “amyloid hotspots” ([7, 24, 25] and references therein). Specific regions proposed in separate studies to drive LC misfolding varied depending on the protein and the experimental approach, underscoring the complexity of amyloidogenic pathways and their modulators. This complexity greatly complicates ongoing efforts to develop therapeutic targets to prevent or decelerate amyloid deposition in this hitherto incurable disease [6].

In the current study, we explored how local LC conformation and dynamics in the native state influence amyloid formation. To this end, we combined biochemical, spectroscopic, and bioinformatics methods with hydrogen-deuterium exchange (HDX) mass spectrometry (MS) to explore 14 patient-derived and engineered LCs and their domain fragments from two closely related κ1 sub-families. Patient-derived proteins were associated with either AL amyloidosis or multiple myeloma (MM), another monoclonal Ig disease wherein most patients do not develop AL [26]. In our study, MM LC was used as a naturally occurring non-AL control. Prior HDX studies using NMR or MS were limited to several LCs from the λ family [21, 25, 27]; the current study extends this approach to the κ family. Our results identified CDR1 and CDR3, which are linked via the internal disulfide, as key factors in κ LC misfolding.

## Methods

### Protein selection, expression, purification and quality control

Proteins related to two germline genes, *IGKVLD-33*01* and *IGKVLD-39*01*, are described in parts 1 and 2, respectively. In Part 1, we explored two patient-based variants, AL 00-131 and MM 96-100 [10], and their common germline LC, GL 33*01 (henceforth termed AL1, MM1 and GL1). Each recombinant protein was generated in two forms, a full-length LC and the separate or stand-alone V_L_ domain (we refer to V_L_ and C_L_ as “domains” in the form of stand-alone constructs and as “regions” in the context of the full-length LC). The separate or stand-alone C_L_ domain, which is common for all LCs in the current study, was also generated. Additionally, three full-length LCs were engineered. One (termed AL1_CDR1_) differed from AL1 by two restorative GL1-like substitutions in CDR1 (G28D, Y32S); another (termed GL1_CDR3_) differed from GL1 by four AL1 substitutions in CDR3 (D92A, N93S, L94V, and P95F); and the third (termed AL1_CDR3_) differed from GL1 by four AL1 substitutions outside CDR3 (D28G, Y32S, S63T, and T74S). In Part 2, we explored four full-length LCs: a patient-based AL 05-009 protein, termed AL2; the corresponding germline protein related to the *IGKVLD-39*01* gene, termed GL2; an engineered protein termed GL2_CDR3_ whose sequence differed from GL2 by four AL2 substitutions in CDR3 (S91C, Y92H, T94I, I96F); and an engineered protein termed AL2_CDR3_ whose sequence differed from GL2 by the five AL2 substitutions outside CDR3 (T20I, S28T, S30R, L46N, A84G). In total, 14 recombinant proteins were generated (Supplementary Table S1).

Proteins were expressed in *E. coli* and purified following published protocols [17]. Recombinant AL LC produced by this method showed similar structural and stability properties as the urinary AL00-131 LC, suggesting that the N-terminal His-tag that was retained to improve solubility did not significantly alter protein structure and stability [17]. Protein purity, sequence, and the lack of post-translational modifications or non-native disulfides were verified by SDS PAGE and mass spectrometry (Table S1, Figure S1). Similar to patient-based proteins, recombinant proteins contained no post-translational modifications. Freshly expressed recombinant proteins in standard buffer (5 mM sodium phosphate, pH 7.4, 15 mM NaCl) were used for further studies.

### Protein analysis

The proteins were explored using bioinformatic, biophysical, and biochemical methods following published protocols [25]. Briefly, amyloid-forming propensity was predicted using the consensus sequence analysis software AmylPred2 [28]. Protein secondary structure and thermal stability were assessed by far-UV circular dichroism (CD) spectroscopy. Proteolytic susceptibility was assessed using limited proteolysis by trypsin, monitored by SDS PAGE. Amyloid formation was monitored by thioflavin T (ThT) binding/fluorescence during protein incubation under amyloid-promoting conditions as follows: 0.3 mg/mL protein in standard buffer at 37 °C with shaking in the presence of 5 μg/mL trypsin (which augments amyloid formation by full-length LCs), 0.5 mM SDS (which augments amyloid formation by LCs and other protein *in vitro*), or 0.02 mg/mL heparin (which is a common amyloid co-factor *in vitro* and *in vivo*) ([25, 29] and references therein). Amyloid aggregates were visualized using transmission electron microscopy (EM) of negatively stained samples. The experimental details of these methods are reported in the Supplementary Appendix.

### Hydrogen deuterium exchange mass spectrometry measurements

HDX MS analyses were performed using methods previously described [25], with modifications, as detailed in the Supplementary Appendix. The recommended summary [30] of HDX MS experimental parameters, proteolytic maps for all proteins, details of replicates acquired, and the numeric values used to create all HDX MS figures are provided in the Supplemental Datafile 1. The HDX MS data have been deposited to the ProteomeXchange Consortium via the PRIDE [31] partner repository with the dataset identifier PXD039682.

## Results

The propensity to form amyloid based on amino acid sequence was predicted using AmylPred2 software. Figure 1 shows the predicted amyloidogenic segments mapped on the LC structure. Despite overall similarities in the amyloid hotspot locations, variant LC sequences showed some differences; *e*.*g*., a major hotspot extending from the β-strand F_V_ (βF_V_) into CDR3 was predicted only in AL1 and AL2 but not in other proteins (Fig. 1A).

### Part 1. Proteins related to κ1 germline gene IGKVLD-33*01

#### Structural stability and amyloid formation by AL1, MM1 and GL1 variants

Global protein conformation and stability were explored by CD spectroscopy and limited proteolysis (Fig. 2). Far-UV CD spectra of AL1, MM1 and GL1 LCs were similar, indicating similar secondary structures consistent with Ig fold (Fig. 2A). The CD melting data showed small but significant differences, with apparent melting temperatures increasing from T_m,app_=48±1 °C in AL1 to 50±1 °C in MM1 and 52±1 °C in GL1 LC. Hence, protein thermodynamic stability increased in order AL1 < MM1 < GL1 (Fig. 2B). Limited proteolysis monitored by SDS PAGE revealed faster degradation of AL1 *vs*. MM1 and GL1 LCs. AL1 LC showed lower initial population of the covalent dimer and faster degradation of the LC monomer (marked LC_2_ and LC_1_ in Fig. 2C) as compared to MM1 and, especially, GL1 LC. In summary, CD and limited proteolysis showed the same rank order of protein stability, AL1 < MM1 < GL1 (Fig. 2B).

**Figure 2.**
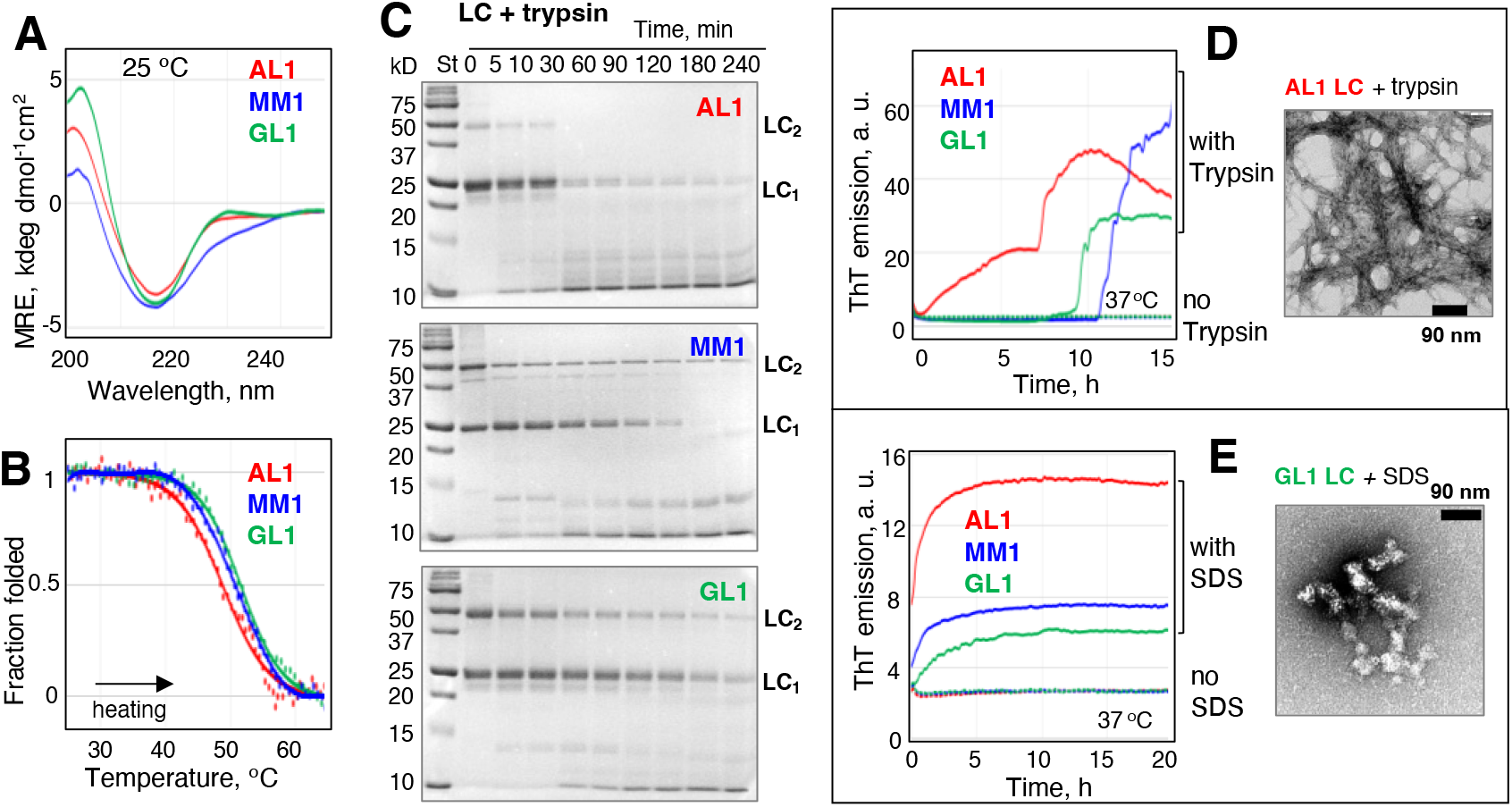
Secondary structure, thermal stability, proteolytic susceptibility and amyloid formation by the patient-derived LCs related to germline gene *IGKVLD-33*01*. In all figures, the data are similarly color-coded: AL1 (red), MM1 (blue), GL1 (green). (**A**) Far-UV CD spectra at 25 °C were recorded using 0.3 mg/mL protein in standard buffer. (**B**) CD melting data were recorded at 206 nm during sample heating at a rate of 0.5 °C/min. (**C**) Limited proteolysis monitored by SDS PAGE. LCs were incubated at 37 °C with trypsin (1:100 enzyme:substrate mg:mg); the incubation times are indicated on the lanes. St - molecular weight standards. Positions corresponding to the migration of the covalent dimers linked via Cys214 (LC_2_) and monomers (LC_1_) are indicated. (**D, E**) Amyloid formation monitored by ThT fluorescence and transmission EM. The proteins were incubated at 37 °C with shaking either with trypsin (D) or SDS (E). ThT emission at 482 nm was used to monitor formation of amyloid-like structure. Representative data for each protein are shown. A representative electron micrograph (negative stain) of AL1 LC after 50 h incubation with trypsin shows fibrillary aggregates. MS analysis of these aggregates showed that they contained largely V_L_ domains.

Next, AL1, MM1, and GL1 LCs were incubated under amyloid-promoting conditions with either trypsin or SDS and formation of amyloid-like structure was continuously monitored by ThT fluorescence. During incubation with trypsin, MM1 and GL1 LCs showed sigmoidal reaction kinetics characteristic of nucleation-growth, while AL1 showed biphasic growth without a lag phase (Fig. 2D). Therefore, amyloid nucleation was much faster in AL1 *vs*. GL1 or MM1 LC. Electron micrographs of the post-incubation samples of AL1 LC with trypsin showed amyloid fibrils (Fig. 2D); MS analysis of these fibrils indicated that they were comprised of V_L_ fragments. During incubation with SDS, which typically eliminates the lag phase and accelerates amyloid formation by LCs ([25] and references therein), AL1 LC showed a greater increase in ThT emission amplitude and a faster growth as compared to either MM1 or GL1 LCs (Fig. 2E). EM of the post-incubation samples showed curvilinear assemblies of spherical aggregates (Fig. 2E).

In summary, the results in Figure 2 show that AL1 LC forms amyloid faster than MM1 or GL1 LCs. Moreover, the rate of amyloid formation in the presence of heparin increased in order GL1 < MM1 < AL1 for both full-length LCs and their V_L_ fragments (Fig. S2). Together, these results are consistent with the observed rank order of thermal and proteolytic stability for these LCs, AL1 < MM1 < GL1 (Fig. 2).

#### HDX MS studies of AL1, MM1, and GL1 proteins

HDX MS was used to explore local protein conformation and dynamics in LC variants. Figure 3A shows deuterium uptake curves (percent deuteration *vs*. exchange time) for full-length LCs. Figure 3B shows an HDX MS data summary as a “skyline plot” (relative deuteration *vs*. residue number) for LCs after 10 min deuteration (HDX MS for all exchange times for LCs and their stand-alone V_L_ and C_L_ domains are shown in Fig. S3).

**Figure 3.**
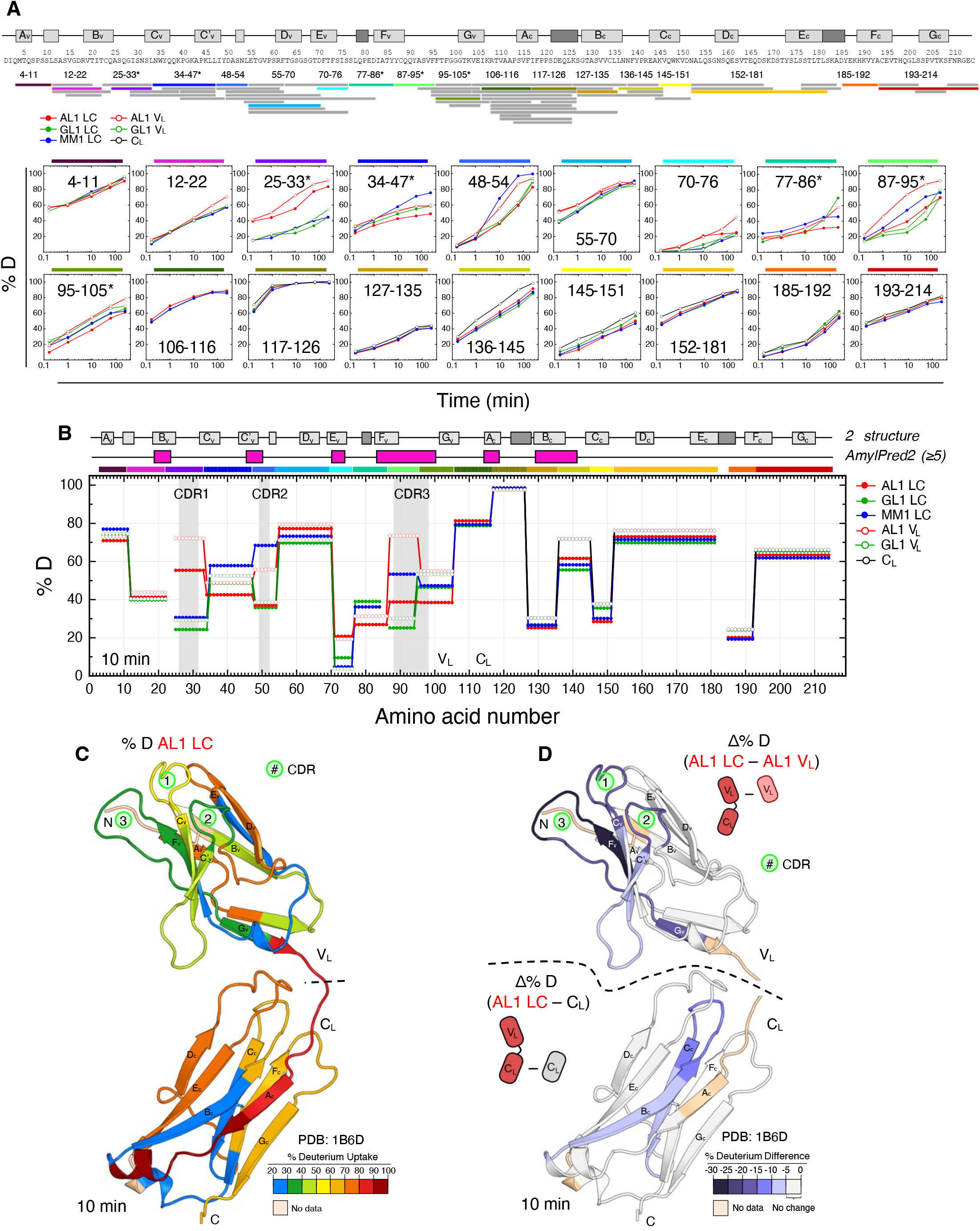
HDX MS analysis of patient-derived proteins related to *IGKVLD-33*01*. **(A)**Peptide coverage map for AL1 LC and deuterium uptake curves for selected peptide fragments (color-coded N- to C-terminus) of AL1, MM1, GL1 full-length LCs (solid symbols) and their stand-alone V_L_ and C_L_ domains (open symbols). All peptides that were followed by HDX MS are indicated with bars below the sequence. The diagram at the top shows primary structure and a linear representation of the native secondary structure in κ1 LC (β-strands in light gray, α-helices in dark gray). **(B)**Skyline plots for full-length LCs and their stand-alone V_L_ and C_L_ domains. Diagrams at the top show native secondary structure and predicted amyloid hotspots that have the AmylPred2 consensus number ≥5 (purple). These data are for the 10 min exchange time for the peptide fragments that were detected in all proteins. Data for all exchange times are shown in Figure S3. **(C)**Percent deuteration values (% D) for AL1 at the 10 min exchange time projected on the κ1 LC crystal structure. See Figure S8 for other exchange times **(D)**Difference in HDX protection between AL1 and AL1 V_L_ (top), and AL1 and C_L_ (bottom) mapped on the crystal structure of κ1 LC, for the 10 min exchange time (see also Fig. S8). Cartoons illustrate the deuterium level subtraction equation.

First, we compared HDX MS results for LC, V_L_ and C_L_ forms of AL1 and GM1. Full-length LCs and their corresponding stand-alone domains showed similar protection in most regions (Fig 3, Fig. S3), suggesting that HDX protection depends mainly on the local conformational dynamics, while the contributions from domain interactions are relatively modest. Nevertheless, several AL1 segments, particularly CDR3, followed by CDR1 and CDR2, showed much higher protection in full-length LC *vs*. V_L_ alone. This is not surprising since, of all flexible CDR loops, CDR3 is closest to the V_L_ – C_L_ cleavage site. GL1 LC showed a similar but smaller trend; moreover, CDRs were much better protected in GL1 V_L_ *vs*. AL1 V_L_. Furthermore, AL1 showed a greatly decreased protection in CDR3 of the stand-alone V_L_ *vs*. LC; this is potentially important, as CDR3 overlaps a major amyloid hotspot in AL1 (Fig. 1). Taken together, these results suggest that if LC cleavage and release of V_L_ generates the protein precursor of amyloid, then amyloid formation by AL1 V_L_ *vs*. GL1 V_L_ can be augmented via two mechanisms: i) increased exposure/dynamics in the CDR3 hotspot of AL1 V_L_ can initiate aggregation; ii) increased dynamics in CDR1 and CDR3 of AL V_L_, which are linked via the nearby C23-C88 disulfide, accelerates polypeptide chain rotation around this disulfide at the rate-limiting step of amyloid formation.

Second, we focused on the common features in HDX profiles of AL1, MM1 and GL1 full-length LCs. Compared to C_L_, much larger variations in protection induced by AL1 or MM1 mutations were observed in V_L_, consistent with its variable nature (Fig. 3). A similar effect was observed in HDX studies of λ6 LCs [25]. Furthermore, for all LCs, V_L_ segments with lowest protection (highest deuteration, most dynamic/exposed) included residues 1-10 (βA_V_ and adjacent segments) followed by 55-72 (C’_V_D_V_ linker, βD_V_, and D_V_F_V_ linker). Conversely, V_L_ segments with highest protection (least deuteration, well-ordered/sequestered) included residues 12-25 in AL1 (12-35 in GL1 and MM1 stretching from A_V_B_V_ linker to βC_V_), and 72-88 (from βE_V_ to βF_V_) (Fig. 3B-D). In C_L_, segments with the lowest protection included A_C_B_C_ linker (around residue 120), while the highly protected segments centered at residues 130 (βB_C_), 150 (βC_C_) and 190 (E_C_-F_C_ linker).

Third, we focused on amyloid hotspots whose decreased protection may help initiate amyloidosis. In all LCs, all amyloid hotspots predicted in V_L_ residues 18-22, 31-37, 45-50, 70-76 and 83-89 (in GL1 and MM1) or 83-96 (in AL1) showed high protection from HDX (Fig. 1). Except for the highly protected βE_V_ residues 70-76, none of these hotspots showed substantially decreased protection in AL1 *vs*. GL1 and MM1 proteins. Therefore, decreased protection in V_L_ hotspots was unlikely to account for the enhanced amyloid formation by the full-length AL1 LC. In C_L_, one of the two predicted amyloid hotspots in βA_C_ residues 114-117 showed low protection, which was similar for AL1, MM1 and GL1 LCs. The other major amyloid hotspot overlapped residues 136-145, which showed slightly lower protection in AL1 *vs*. MM1 and GL1 (Fig. 3). Since the overall protection in this segment was relatively high, it was unlikely to account for the enhanced amyloid formation by AL1 LC.

Fourth, we focused on residues 105-125 from the V_L_-C_L_ linker that are important for LC domain orientation, structural integrity, and amyloidogenicity ([23] and references therein). High proteolytic susceptibility of this linker can promote the generation of V_L_ fragments, such as those found in *ex vivo* AL deposits [18]. This linker showed relatively low protection from HDX that was similar for all LCs, with only a slight decrease seen in AL1 *vs*. MM1 and GL1 LCs (Fig. 3), suggesting comparable proteolytic susceptibility in AL *vs*. non-AL LCs. In summary, neither sensitive amyloid hotspots nor the inter-domain linker showed a large decrease in HDX protection for AL1 *vs*. non-AL LCs. Therefore, the main amyloid-contributing factor emerging from our studies of full-length LCs is decreased overall stability of AL1 *vs*. MM1 and GL1 LC.

Next, to compare mutational effects on the local *vs*. global protein stability, we examined differences in HDX protection for all segments. The largest decrease in AL1 protection at all exchange times (Fig. 3, Fig. S3) was observed in residues 25-35 that overlap CDR1 and contain two substitutions, D28G and Y32S, differentiating AL1 from GL1 (henceforth termed AL1 substitutions). This local destabilization near the mutation site correlated with the changes in the overall stability of LC, AL1 < MM1 < GL1, and could potentially contribute to the overall destabilization of AL1 *vs*. GL1 LC. AL1 substitutions in other regions, including S63T in βD_v_, T74S in βE_v_, and D92A, N93S, L94V, P95F in CDR3, caused much smaller changes in local HDX protection (Fig. 3, Fig. S3), which did not correlate with global protein stability. Taken together, our HDX MS and stability studies suggest that AL1 substitutions D28G and Y32S cause local destabilization in CDR1.

#### Probing the role of CDR1 by engineered substitutions

To test the role of CDR1, we explored an engineered LC, termed AL1_CDR1_, which differed from AL1 in two restorative substitutions, G28D and S32Y, in CDR1 (Fig. 4). A different far-UV CD spectrum suggested altered conformation in AL1_CDR1_ LC compared to either AL1 or GL1 LCs (Fig. 4A). CD melting data of AL1_CDR1_ LC showed T_m, app_ = 46.5±1 °C, a small but significant decrease compared to T_m,app_ = 48.0 ±1 °C in AL1, suggesting the rank order of LC thermal stability AL_CRD1_ < AL1 < GL1 (Fig. 4B). These unexpected results indicated context-dependent interactions among the mutation sites in CDR1 and other regions. Limited proteolysis showed comparable proteolytic susceptibility in AL1 and AL1_CDR1_ LC (Fig. 4C). During amyloid formation in the presence of trypsin, SDS or heparin, AL1_CDR1_ LC showed greater changes in ThT emission amplitude as compared to either AL1 or GL1 LC (Fig. 4D, E), suggesting structural differences in amyloids formed by these proteins.

**Figure 4.**
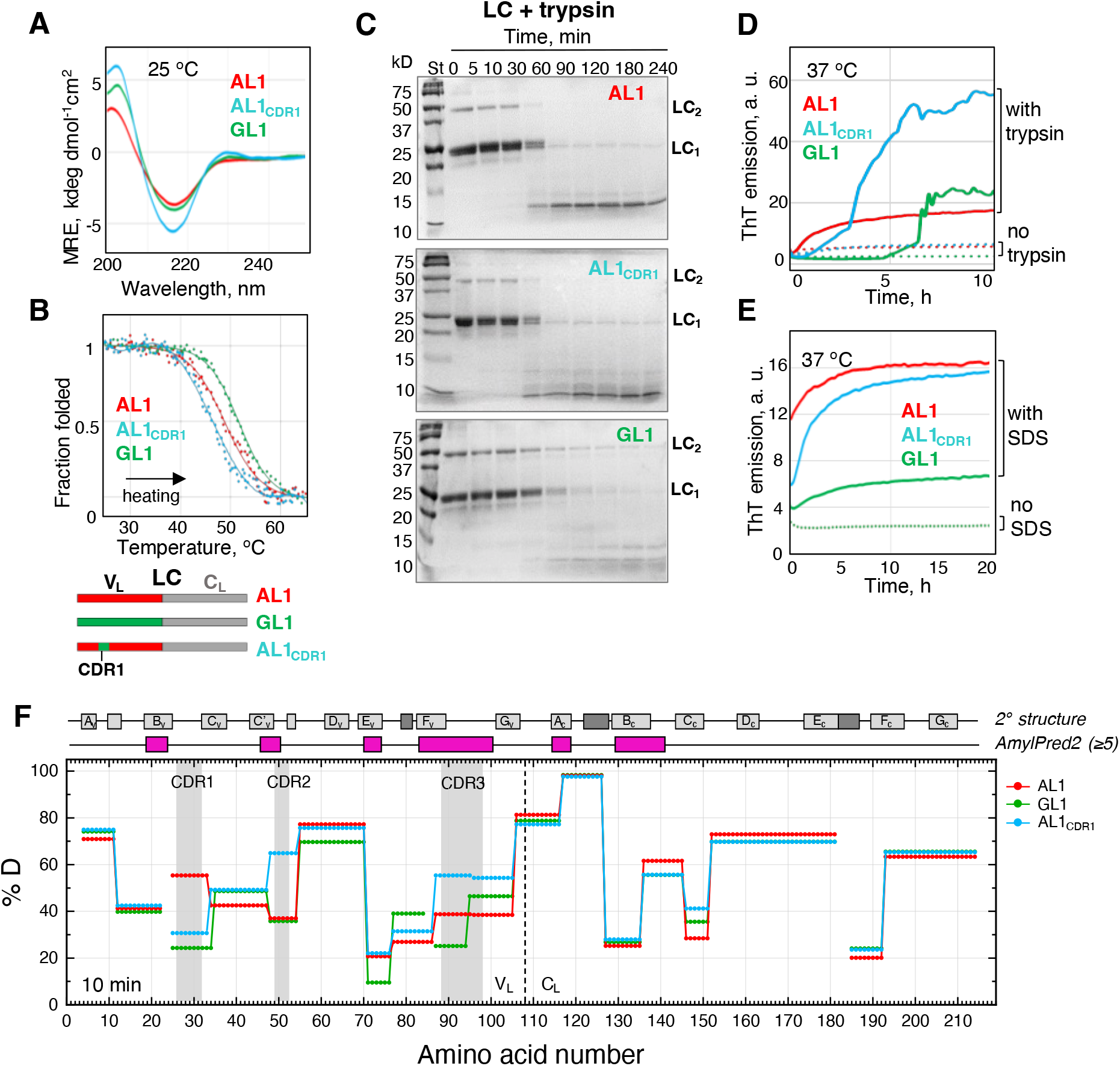
Effects of mutations in CDR1 on the secondary structure, thermal stability, amyloid formation and local dynamics in AL1 LC. Data for the engineered AL1_CDR1_ LC (in teal) are compared to those of AL1 and GL1 LCs. Far-UV CD spectra (**A**), CD melting data (**B**), limited proteolysis by trypsin monitored by SDS PAGE (**C**), the time course of amyloid formation in the presence of trypsin (**E**) or SDS (**D**), and the skyline plots summarizing the HDX MS results for the 10 min exchange time (**F**) were obtained as described in Figure 3 and in the Supplementary Appendix. Data at all HDX MS points are found in Fig. S4.

HDX protection in AL1 LC and AL1_CDR1_ LC was similar but not identical. As expected, the largest differences were observed in CDR1 that at all labeling time points showed much better protection in AL1_CDR1_ vs. AL1 (Fig. 4F, Fig.S4). This supports our conjecture that AL1 mutations D28G and Y32S in CDR1 are locally destabilizing. The mutations also led to altered HDX protection in other regions; *e*.*g*., CDR2 and CDR3 showed lower protection in AL1_CDR1_ *vs*. either AL1 or GL1, which could relate to the lower thermal stability of AL1_CDR1_. Furthermore, the mutations induced small alterations in protection of C_L_ segments, *e*.*g*., near residues 140 and 150 (Fig. 4F). This illustrates propagation of mutation-induced dynamic changes from CDR1 to distant sites. A similar effect was inferred from the analyses of AL1 *vs*. GL1 LC (Fig. 3) and λ6 LCs [25].

Collectively, our results show that AL1 substitutions in CDR1 are locally destabilizing (Fig. 4F). However, GL1-like replacements in CDR1 of AL1_CDR1_ LC did not restore thermal stability (Fig. 4B). Moreover, AL1_CDR1_ LC showed altered conformations (Fig. 4A) and kinetics of amyloid formation as compared to AL1 and GL1 (Fig. 4D), suggesting non-additive effects of AL1 substitutions within and outside of CDR1. Therefore, interactions between mutation sites in CDR1 and other sites contribute to the decreased stability of AL1 LC and its amyloidogenic properties. Spatial proximity of CDR1 to CDR3, which harbors four out of eight AL1 substitutions (Fig. 1), suggests that these other sites probably include CDR3.

#### Probing the role of CDR3 by engineered substitutions

To determine how AL1 substitutions within and outside of CDR3 influence protein stability, two engineered LCs were explored: GL1_CDR3_, containing AL1 replacements only within CDR3 (D92A, N93S, L94V, and P95F); and AL1_CDR3_, containing AL1 replacements outside of CDR3 (D28G, Y32S, S63T, and T74S). The effects of these replacements on protein stability, amyloid formation, and local conformational dynamics are shown in Figures 5, 6.

**Figure 5.**
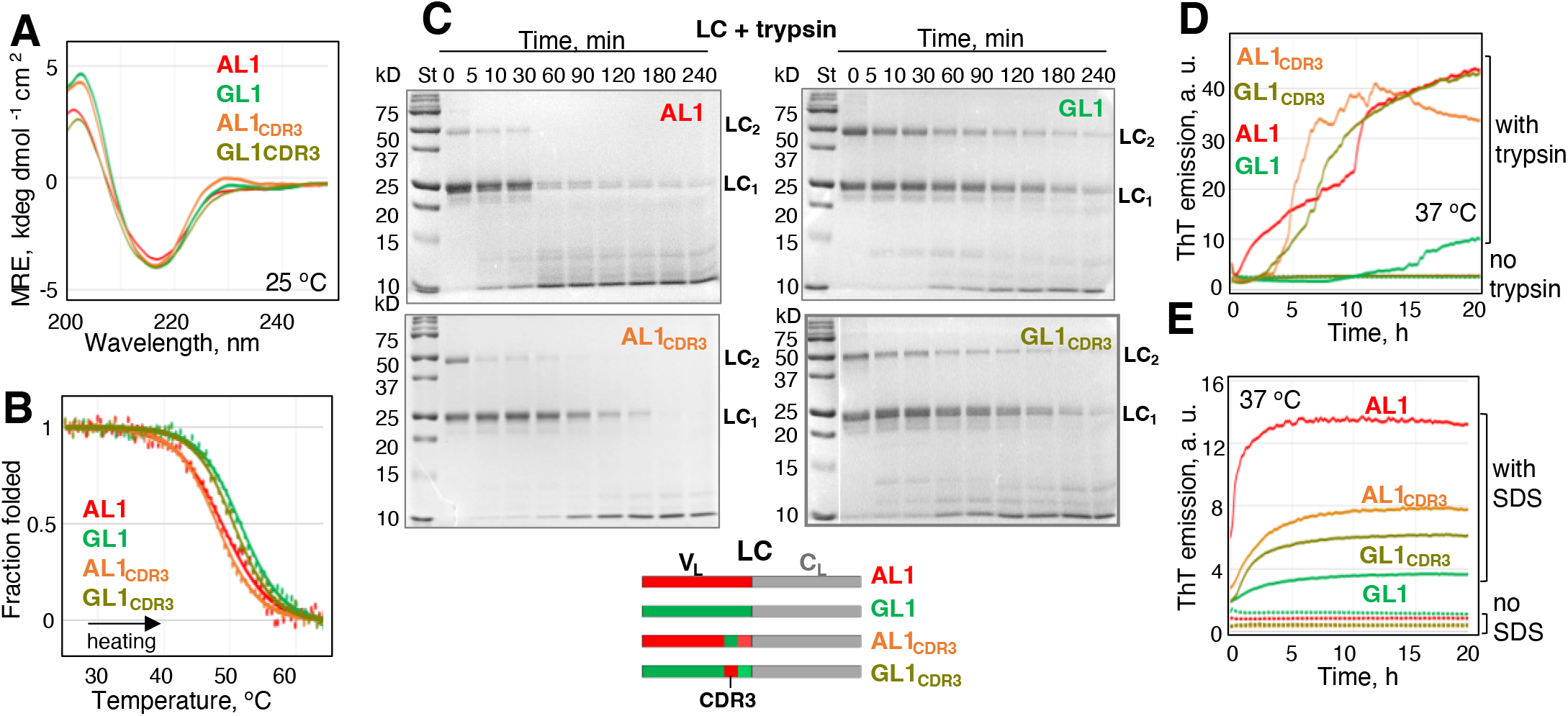
Secondary structure, thermal stability, limited proteolysis and amyloid formation by engineered LCs, GL1_CDR3_ (olive) and AL1_CDR3_ (orange), compared with GL1 (green) and AL1 (red). Schematics at the bottom illustrates the primary structures of the engineered proteins. Far-UV CD spectra (**A**), CD melting data (**B**), limited proteolysis monitored by SDS PAGE (**C**), and the time course of amyloid formation monitored by ThT fluorescence in the presence of trypsin (**D**) or SDS (**E**). Experimental conditions are described in Methods and in Figure 2 legend. Data for amyloid formation in the presence of heparin are found in Fig. S5.

**Figure 6.**
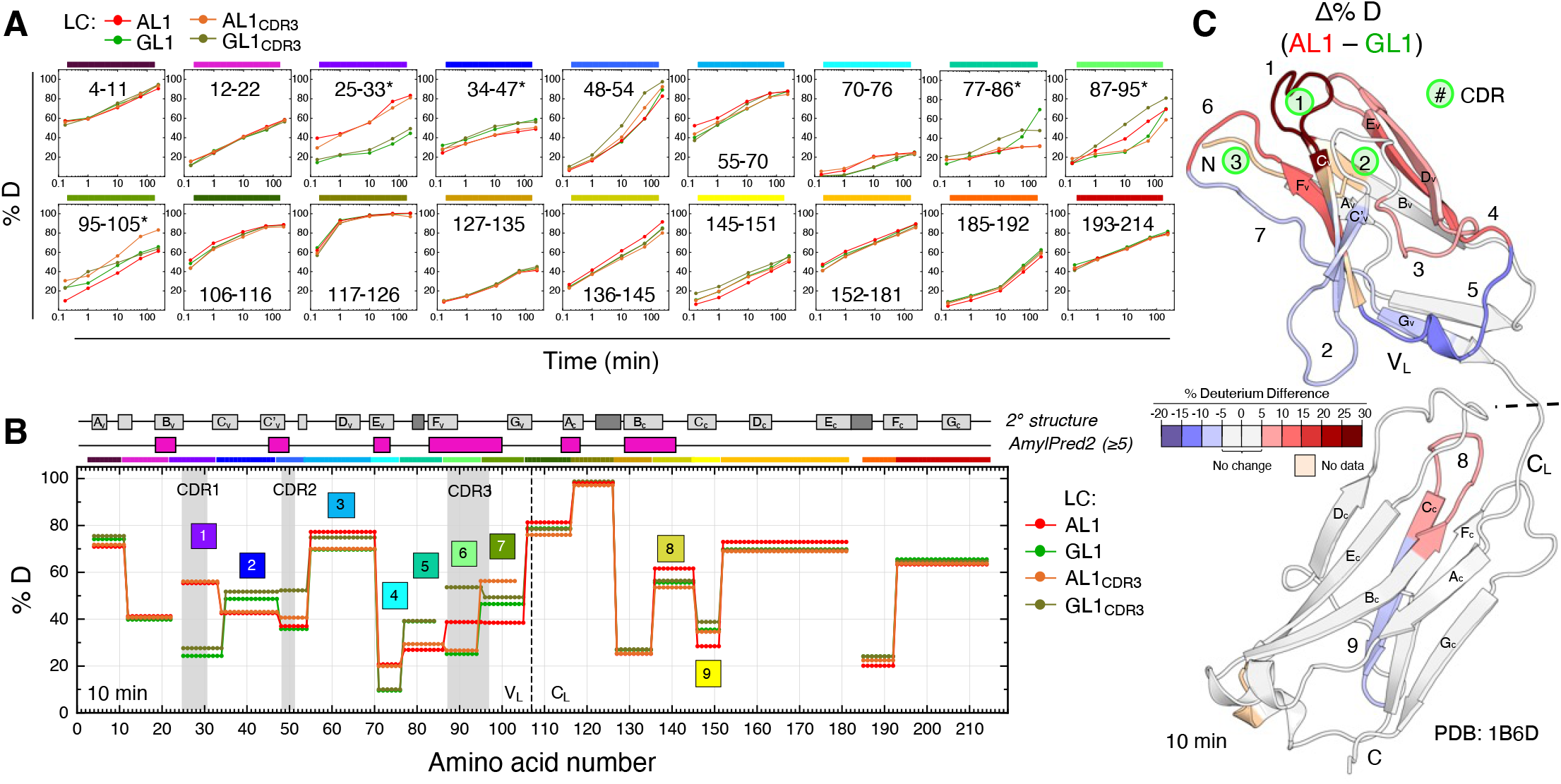
HDX MS analysis of AL1, GL1, AL1_CDR3_ and GL1_CDR3_. (**A**) Uptake curves for selected peptide fragments. Fragments are color-coded according to Figure 3 schematics; residue numbers are indicated. (**B**) Skyline plot for selected peptides after 10 min exchange. (**C**) Difference in HDX after 10 min exchange of AL1 and GL1 LCs, (AL1 – GL1), mapped on the crystal structure of κ1 LC. For other exchange times see Figs. S4 and S8.

Far-UV CD spectra of AL1, GL1, AL1_CDR3_ and GL1_CDR3_ LCs suggested similar overall secondary structures (Fig. 5A). The rank order of thermal stability determined by CD, AL1_CDR3_ ≤ AL1 < GL1 _CDR3_ < GL1, was consistent with proteolytic stability determined by tryptic digestion, AL1 ≅ AL1_CDR3_ < GL1_CDR3_ ≅ GL1 (Fig. 5B, C). This order suggested that reduced overall stability of AL1 *vs*. GL1 LC stems largely from substitutions outside of CDR3 (Fig. 1). Overall stability correlated directly with the fraction of covalent LC dimers, which in AL1 and AL1_CDR3_ was lower than in GL1 and GL1_CDR3_ as observed by SDS PAGE (lane 0, Fig. 5C). The rate of amyloid formation in the presence of SDS increased in order GL1 < GL1_CDR3_ ≅ AL1_CDR3_ < AL (Fig. 5E), which was the inverse of protein stability order and was influenced by mutations both within and outside of CDR3. Amyloid formation with trypsin showed that the length of the lag phase increased from 0 h to ∼15 h in order AL1 < AL1_CDR3_ ≅ GL1_CDR3_ < GL (Fig. 5D), which again was inverse to protein stability ranking. Taken together, the results in Fig. 5 indicate that decreased stability of AL1 *vs*. GL1 LC stems mainly from the substitutions outside CDR3, while enhanced amyloid formation by AL1 *vs*. GL1 LC results from the combined effects of substitutions within and outside of CDR3.

Interestingly, the amyloid formation kinetics in the presence of heparin (Fig. S5), a common amyloid cofactor, showed a different rank order, with the fastest amyloid formation observed in GL1_CDR3_ LC and the slowest in GL LC, suggesting that AL1 mutations altered amyloid-heparin interactions.

HDX MS consistently showed that the largest differences induced by AL1 mutations were located in CDR1; segment 25-34 showed much lower protection at all exchange times in AL1 and AL1_CDR3_ *vs*. GL1 and GL1_CDR3_ LCs (Fig. 6, Fig. S4). This local destabilization resulted from D28G and Y32S substitutions, as indicated by HDX analysis of AL1_CDR1_ (Fig. 4). HDX protection in CDR3 showed mixed results; in residues 87-94, AL1 showed similar or lower protection than GL1 at all times, GL1 and AL1_CDR3_ showed similar protection, while GL1_CDR3_ showed lower protection (higher exchange) than any other protein (Fig. 6 uptake curves). This indicates context-dependent mutational effects: AL1 substitutions in CDR3, which caused modest local destabilization in the presence of other AL1 mutations (compare AL1 with GL1), became more destabilizing in the context of GL1 (compare GL1_CDR3_ with GL1), implying that there must be interactions among mutation sites within and outside of CDR3.

#### Part 1 summary

We used far-UV CD spectroscopy, limited proteolysis, *in vitro* amyloid formation, and HDX MS to explore patient-derived proteins (including AL1, MM1 and GL1 LCs and their stand-alone V_L_ and C_L_ domains) and engineered LCs (AL1_CDR1,_ AL1_CDR3_ and GL1_CDR3_) related to κ1-family germline gene *IGKVLD-33*01*. The results in Figures 2-6 and S2-S5 revealed several important findings. First, for patient-derived proteins, LC stability increased in order AL1 < MM1 < GL1 and for engineered *vs*. patient-derived proteins, the stability increased in order AL1_CDR1_, AL1_CDR3_ ≤ AL1< GL1_CDR3_ < GL1; this order was approximately inversely related to the rate of amyloid formation in the presence of trypsin or SDS. In addition, reduced stability of AL1 *vs*. GL1 LC stemmed mainly from the four AL1 substitutions outside CDR3, two of which were located within CDR1, while the enhanced amyloid formation by AL1 *vs*. GL1 LC stemmed from combined effects of substitutions within and outside of CDR3. The latter suggests that overall destabilization of full-length LC does not fully explain enhanced amyloid formation by AL1 *vs*. GL1 LC.

One potential trigger of amyloid formation emerging from our results is the increased amyloidogenic sequence propensity of the hotspot overlapping CDR3 (Fig. 1) combined with its increased dynamics/exposure observed in the stand-alone V_L_ domains of AL1 *vs*. GL1 (Fig. 4). However, the causal role of the misfolding and aggregation of V_L_ fragments in AL amyloidosis remains to be established [18]. Unlike stand-alone V_L_, full-length AL1 LC showed no major local destabilization either in the amyloid hotspots or in the inter-domain linker (HDX in Figs. 3, 4, 6). Therefore, mutational effects on the native conformation of LCs do not fully explain why AL1 LC is more amyloidogenic than GL1 LC. This suggests the importance of other factors, such as mutational effects on the misfolded protein state (see Discussion).

#### Part 2. LCs related to κ1 germline gene IGKVLD-39*01

##### Structural stability and amyloid formation by AL2, GL2, AL2_CDR3_ and GL2_CDR3_ variants

In Part 2, we used similar techniques as in Part 1 to explore LCs from two closely related κ1 subfamilies, but the results were surprisingly different. First, global protein conformation and stability of GL2, GL2_CDR3_ AL2, and AL2_CDR3_ LCs were explored by CD spectroscopy and limited proteolysis. Compared to other proteins, AL2_CDR3_ LC showed a significantly altered far-UV CD spectrum, indicating non-native secondary structure (Fig. 7A) along with enhanced high-order aggregation observed by SDS PAGE (Fig. 7C, lane 0). As in Part 1, covalent dimerization was decreased in AL2 LC *vs*. GL2 LC (Fig 7C, lanes 0). Nevertheless, CD melting data of AL2 and GL2 LCs overlapped, indicating similar overall thermodynamic stability with T_m,app_ = 51±1 °C; AL2_CDR3_ showed a large low-temperature shift to T_m,app_ =44±1.5 °C, while GL2_CDR3_ showed a large high-temperature shift to T_m, app_=58±1 °C (Fig. 7B). Hence, thermal stability increased in order AL2_CDR3_ << AL2 ≅ GL2 < GL2_CDR3_. This unexpected finding was supported by limited proteolysis showing that proteolytic stability increased in order AL2_CDR3_ << AL2 ≅ GL2 < GL2_CDR3_ (Fig. 7C). These results indicate that, compared to GL2-like CRD3 which is destabilizing (in AL2_CDR3_ and GL2 LCs), AL2-like CDR3 is stabilizing both in AL2 and in GL2 contexts (in AL2 and GL2_CDR3_ LCs).

**Figure 7.**
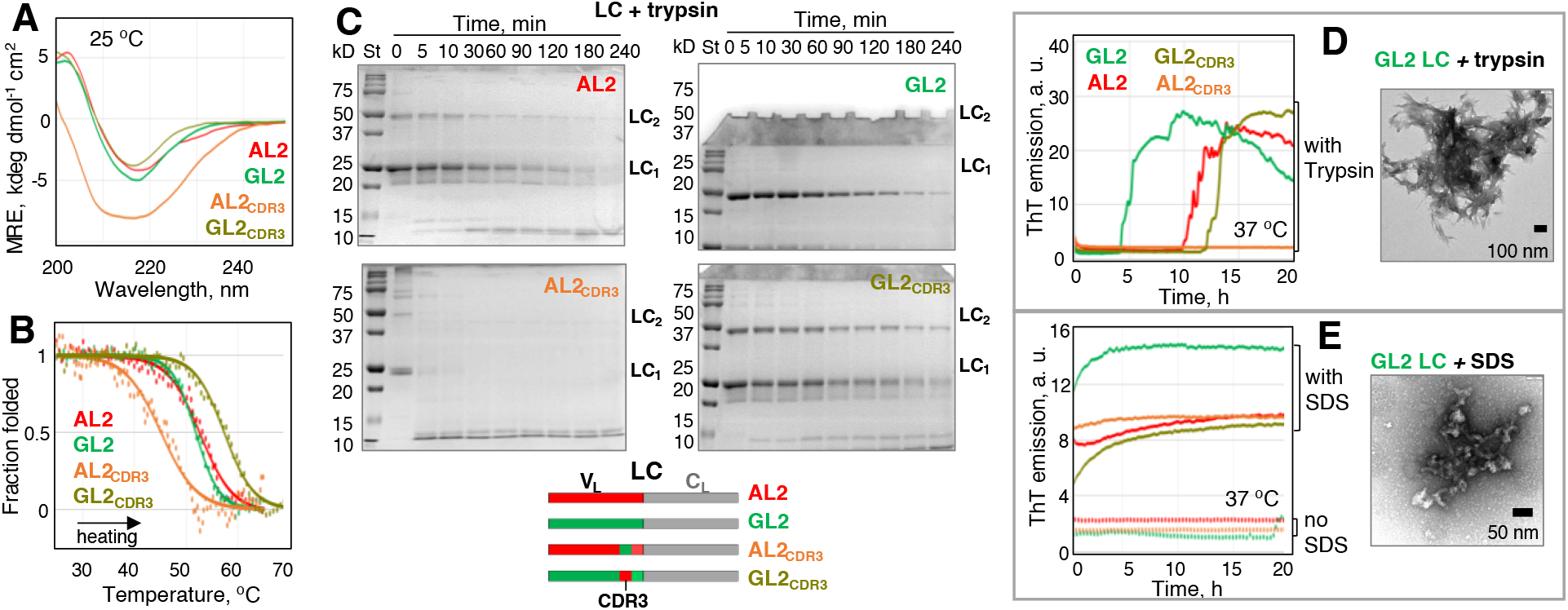
Secondary structure, thermal stability, limited proteolysis and amyloid formation by the patient-derived and engineered LCs related to the germline gene *IGKVLD-39*01*: AL2 (red), GL2 (green), GL2_CDR3_ (olive) and AL2_CDR3_ (orange). Schematics at the bottom illustrates the primary structures of the engineered proteins. Far-UV CD spectra (**A**), CD melting data (**B**), limited proteolysis monitored by SDS PAGE (**C**), and the time course of amyloid formation monitored by ThT fluorescence in the presence of trypsin (**D**) or SDS (**E**) are shown, together with electron micrographs (negative stain) of the samples after 50 h incubation under amyloid-promoting conditions with either trypsin or SDS (**D, E**). Experimental conditions are described in Methods and in Figure 2 legend. Data for amyloid formation in the presence of heparin are found in Fig. S6.

Next, the proteins were incubated under amyloid-promoting conditions with trypsin. AL2_CDR3_ LC, which was least stable, was rapidly digested and no amyloid formation was detected; other LCs formed amyloid that was detected by ThT fluorescence and EM (Fig. 7D). AL2 consistently showed a ∼10 h lag phase in amyloid formation, while GL2 and GL2_CDR3_ showed large variations in the lag phase that was typically shorter for GL2 *vs*. AL2 LC (Fig. 7D). Therefore, in the presence of trypsin AL2 LC showed comparable or slower amyloid nucleation compared to GL2 LC. Furthermore, in the presence of SDS, AL2 LC showed slower amyloid formation *vs*. GL2 LC (Fig. 7E) and with heparin, the rate of amyloid formation by AL2 and GL2 LCs was comparable (Fig. S6).

##### HDX MS studies of AL2, GL2, AL2_CDR3_ and GL2_CDR3_ LCs

As in Part 1, HDX MS was performed on the proteins in Part 2. Figure 8 shows deuterium uptake curves, skyline plots after 10 min exchange (see Fig. S7 for skyline plots for all HDX MS time points), and a summary of differences in exchange plotted on the crystal structure. HDX MS revealed a number of major local differences in V_L_ regions. Compared to other LCs, AL2_CDR3_ showed greatly decreased protection across most V_L_ segments (except for the N-terminal ∼22 residues and a short helix in residues 79-85, perhaps involved in aggregation). These results are consistent with large spectral changes in far-UV CD and greatly decreased stability of AL2_CDR3_ LC revealed by CD and limited proteolysis (Fig. 7). The decreased stability of AL2_CDR3_ LC was limited to V_L_ while C_L_ remained stable, as indicated by HDX (Fig. 8). In addition, two segments, 12-22 and 79-105, showed unexpectedly higher protection in AL2 *vs*. GL2 LC. These segments are adjacent to or overlap the Cys23-Cys88 disulfide. This finding suggests that decreased dynamics of the polypeptide chain near the disulfide may contribute to delayed amyloid formation observed in AL2 *vs*. GL2 LC in the presence of trypsin (Figs. 7C,D). Moreover, residues 25-32 overlapping CDR1 showed large differences in HDX protection that increased in order AL2_CDR3_ < AL2 < GL2 < GL2_CDR3_, mirroring the rank order of protein stability observed by CD and limited proteolysis (Fig. 7). Furthermore, residue segment 62-73 showed decreased protection in AL2 *vs*. GL2; this may contribute to amyloid formation by AL2 LC, as segment 62-73 partially overlaps a major amyloid hotspot in βE_V_. Lastly, residue segment 87-105, which overlaps CDR3, showed large differences in protection that increased in order AL2_CDR3_ < GL2 < GL2_CDR3_ ≅ AL2. Consequently, AL2-like CDR3 shows substantial local protection in both AL2 and GL2 contexts (in AL2 and GL2_CDR3_); this protection greatly decreased for GL2-like CDR3 (in GL2 and AL2_CDR3_). This result is in strong agreement with CD and limited proteolysis data indicating a stabilizing role of AL-like *vs*. GL-like CDR3 (Fig. 7).

**Figure 8.**
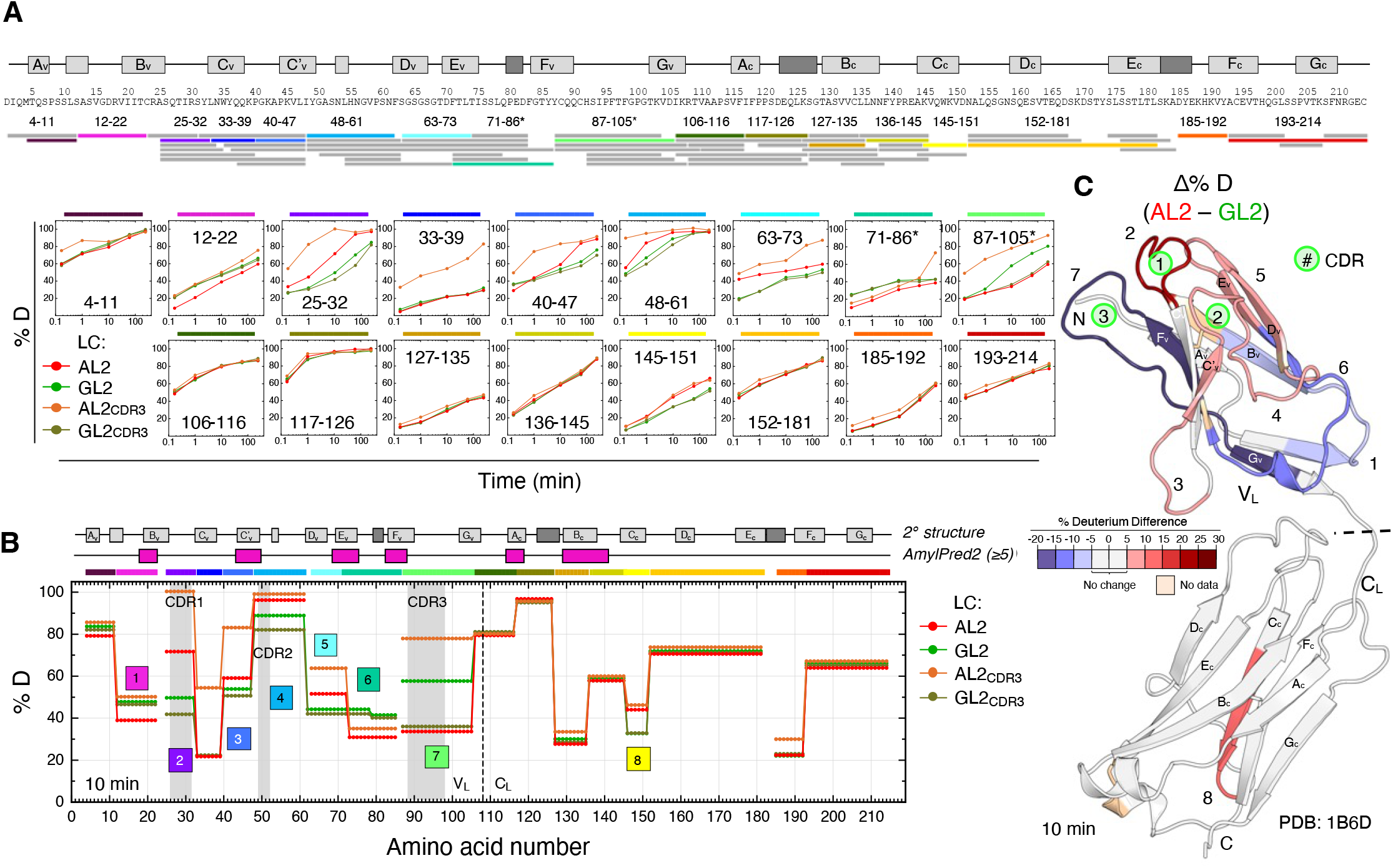
HDX MS of AL2, GL2, GL2_CDR3_ and AL2_CDR3_ LCs. (**A**) Peptide coverage map for AL2 LC and uptake curves for selected peptide fragments. (**B**) Skyline plots after 10 min exchange. (**C**) Difference in HDX after 10 min exchange of AL2 and GL2 LCs, (AL2 – GL2), mapped on the crystal structure of κ1 LC. See also Figs. S7 and S8.

In the J-region and downstream linker segment (residues 106-126), the protection from HDX and resilience to tryptic digestion was comparable for AL2 and GL2 LC (Figs. 7, 8). Therefore, as in Part 1, an enhanced initial cut in the inter-domain linker to generate misfolding-prone V_L_ fragments is unlikely to contribute to the differences in amyloid formation by AL2 *vs*. GL2 LC.

All LCs explored in the current study showed only small variations in HDX protection of C_L_. Some of the largest differences in C_L_ were detected in segment 145-154 encompassing βC_C_, with AL2 and AL2_CDR3_ showing lower protection *vs*. GL2 and GL2_CDR3_ at longer exchange times (Fig. 8). Since βC_C_ is not involved in any known intra- or inter-domain interactions in LC monomer or dimer, these HDX alterations probably reflect propagation of dynamic changes from the mutation sites in V_L_ to a distant βC_C_ site in C_L_.

##### Part 2 summary and comparison with Part 1

The analyses of patient-derived (AL2, GL2) and engineered (AL2_CDR3_ and GL2_CDR3_) LCs related to κ1 germline gene *IGKVLD-39*01* can be summarized as follows. First, unlike Part 1 (proteins related to germline gene *IGKVLD-33*01*), engineered LCs from Part 2 showed major changes in stability that increased in order AL2_CDR3_<< AL2 ≅ GL2< GL2_CDR3_. Second, this order was not inversely related to the rate of amyloid formation. Unexpectedly, compared to GL2, AL2 LC showed comparable or slower amyloid formation. We posit that high protection from deuteration observed by HDX MS in AL2 residue segments 12-22 and 87-105 comes from slow polypeptide chain movement around the conserved internal disulfide, Cys23-Cys88. Third, AL2 substitutions in CDR3 stabilized the protein and counteracted the combined destabilizing effects of AL2 substitutions outside CDR3; three out of five such substitutions were located within or near CDR1 (Fig. 1). Fourth, compared to GL2, AL2 showed decreased protection in parts of two major amyloid hotspots overlapping βC’_V_ and βE_V_ (Fig. 8), raising the possibility that decreased protection of these hotspots initiates amyloid formation by AL2 LC. In summary, decreased protection in amyloid hotspots βC’_V_ and βE_V_ may initiate an early step of aggregation in AL2, while increased protection in segments encompassing the conserved internal disulfide C23-C88 (Fig. 8) helps explain the slower amyloid formation in AL2 observed in the presence of trypsin or SDS (Fig. 7D, E).

## Discussion

The current study strived to link physicochemical and amyloid-forming properties of patient-derived κ1 LCs. Proteins from two closely related germline genes showed unexpected differences: AL1 LC was less stable and formed amyloid faster than GL1 LC (Fig. 5), whereas AL2 LC showed similar stability (thermal and proteolytic) and formed amyloid at a comparable or slower rate than GL2 LC (Fig. 7). Therefore, compared to their non-AL counterparts, AL LCs do not necessarily have decreased overall stability, nor do they necessarily show faster amyloid formation *in vitro* (Fig. 7). Consequently, additional factors must contribute to amyloid formation by AL LCs. These factors may include mutational effects on the local conformational dynamics in specific regions that can destabilize the native structure, stabilize the misfolded structure, and/or alter its interactions with amyloid modulators (summarized below). The interplay between CDR1 and CDR3, which are linked via the conserved disulfide and harbor the majority of AL mutations (6 out of 8 in AL1 and 7 out of 9 in AL2) (Fig. 1), emerges as a major player in amyloid formation.

### Destabilization of the native structure does not fully explain amyloid formation by AL1 LC

Decreased thermal and proteolytic stability likely contributes to enhanced amyloid formation by AL1 *vs*. GL1 LC (Fig. 2). Analysis of patient-based and engineered LCs (Figs. 4-6) suggests that this destabilization stems mainly from the combined effects of four AL1 substitutions outside CDR3: D28G and Y32S in CDR1, S63T in βD_v_, and T74S in βE_v_. What is the structural basis for this destabilization? It is likely that a D28G substitution in the loop causes entropic destabilization in AL1. Furthermore, in the native structure of a closely related κLC (PDB ID: 1B6D, 98% sequence identity to AL1 LC), the Y32 side chain is solvent-exposed and can form an aromatic interaction with the conserved Y91 from CDR3; elimination of this interaction by a Y32S substitution in AL1 is potentially destabilizing (Fig. 9A, B). Lastly, prior studies of λ6 LCs showed that single conservative point substitutions such as N32T at an exposed site in CDR1 can decrease protein stability and promote fibrillogenesis [25, 27]. In the current study, the effects of conservative AL1 substitutions S63T and T74S in solvent-exposed locations outside CDR3, which are difficult to predict, may also contribute to the decreased overall stability of AL1 and AL1_CDR3_ *vs*. GL1 GL_CDR3_ LC (Fig. 5).

**Figure 9.**
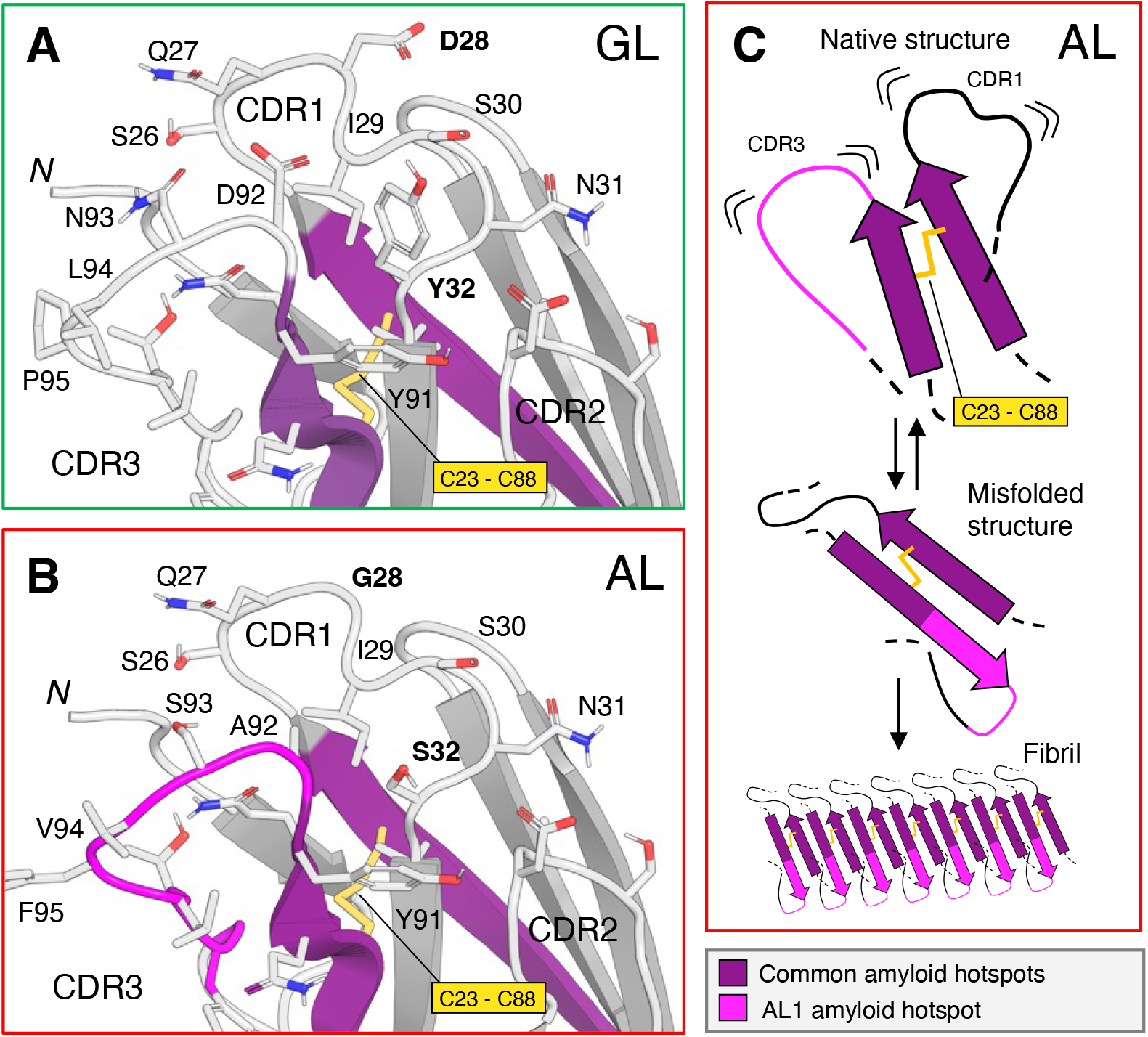
Interactions involving CDR1 linked to CDR3 and the role of AL1 replacements in promoting protein misfolding. (**A, B**) Crystal structure of a related κ1 LC zoomed in on CDRs. Side chains that differ between GL1 and AL1 are shown in (**A**) and (**B**). In GL1, Y32 can interact with the conserved Y91 edge to face to stabilize native CDR1 – CDR3 packing; in AL1, Y32S replacement abrogates this stabilizing interaction. D28G substitution in the loop will likely cause entropic destabilization in AL1. (**C**) Schematics illustrating how increased local flexibility and increased amyloid-forming sequence propensity due to AL1 mutations in CDR1 and CDR3 promote amyloid formation. Increased flexibility accelerates main chain rotation around the C23-C88 disulfide from the parallel (in native V_L_) to the antiparallel orientation (in amyloid). In the antiparallel orientation, the extended hotspot in CDR3 of AL1 (magenta) packs against its counterpart from CDR1, thereby stabilizing amyloid. Amyloid hot spots near CDR1 and CDR3, which are predicted in both AL1 and GL1 (purple) or only in AL1 but not in GL1 (magenta), are colored.

Decreased stability of the native structure does not fully explain enhanced amyloid formation by AL1 *vs*. MM1 or GL1 LC. Therefore, factors other than overall destabilization must contribute to AL1 LC misfolding. Although local destabilization in key regions may be one of such factors, HDX MS analysis of full-length LCs did not show increased dynamics/exposure in either amyloid hotspots or the proteolysis-prone inter-domain linker region of AL1 *vs*. GL1 LC (Fig. 6). This compels us to propose that AL1 mutations exert their effects through other mechanisms, such as altered structures and dynamics of key misfolded states.

### Stabilization of the misfolded state may augment amyloid formation by AL1 and AL2 LCs

Although no atomic structures of κ LC amyloids have been reported to-date, all currently available fibril structures of λ LCs show that amyloid formation involves a 180° rotation around the conserved internal disulfide [9, 13-15]. Therefore, disulfide-linked strands βB_V_ and βF_V_, which are parallel in the native κ LCs crystal structure, are expected to become antiparallel in amyloid (Fig. 9C). In AL1 and AL2 LCs of the current study, βB_V_ and βF_V_ overlap two amyloid hotspots adjacent to CDR1 and CDR3 (Fig 1A). Mutations near these CDRs that enhance the packing of these juxtaposed amyloid hotspots in the fibril core are expected to stabilize amyloid. Figures 1B and 9C illustrate how AL1 mutations, that increase the hydrophobicity of CDR3 and extend its amyloid hotspot C-terminally, may enhance the antiparallel packing against the CDR1 hotspot and thereby stabilize amyloid. Similarly, AL2 shows a modest increase in the amyloid-forming sequence propensity of the hotspots overlapping CDR1 and CDR3 (Fig. 1), which may enhance their juxtaposed antiparallel packing in amyloid. Since CDR3 is the most variable region in κ LC, followed by CDR1 and CDR2, this mechanism may extend to other mutations in or near CDR3 and CDR1 that stabilize the amyloid structure.

Taken together, our studies of AL1 LC suggest that its enhanced amyloid formation stems from decreased overall stability of its native structure (Fig. 2A, C, D) perhaps combined with stabilization of its misfolded state, wherein the antiparallel orientation of the disulfide-linked polypeptide chain segments could be stabilized by mutations in CDR1 and CDR3 (Fig. 1B, Fig. 9C). The latter mechanism may apply to AL2 LC and, potentially, other LCs with mutations in these CDRs. Additionally, decreased protection in a CDR3 hotspot observed in the stand-alone AL1 V_L_ (Fig. 4), combined with increased amyloidogenic sequence propensity of this CDR3 in AL1 (Fig. 1), may potentially contribute to amyloidosis.

### Molecular basis for the unusual behavior of AL2 LC

AL2 and GL2 LCs showed similar stability (Fig. 7), supporting the notion that decreased overall stability of a globular protein is neither necessary nor sufficient for amyloid formation [17, 19]. Analysis of engineered LCs showed that AL2 mutations in CDR3 were stabilizing both locally and globally, while those outside CDR3 were destabilizing (Figs. 7, 8); their combined effects balanced out, leading to similar overall stability of AL2 and GL2. Surprisingly, compared to GL2, AL2 LC showed delayed amyloid formation in the presence of trypsin and lower ThT emission intensity in the presence of SDS (Fig. 7D, E). HDX MS helped explain this observation; segments 12-22 and 73-105 showed increased protection in AL2 *vs*. GL2 (Fig. 8), which is expected to slow the polypeptide chain rotation around the C23-C88 disulfide and thereby decelerate amyloid formation.

Our HDX MS results suggest that amyloid formation by AL2 LC is initiated, in part, by decreased protection of its amyloid hotspots overlapping βC’_V_ and βE_V_ (Fig. 8). Consistent with this conjecture, studies from Eisenberg’s group proposed that the major hotspot in βE_V_ drives amyloid formation in other LCs [24]. Furthermore, as in AL1, AL2 mutations in and near CDR1 (T20I, G28T, and S30R) enhance amyloidogenic sequence propensity in βB_V_ and βC_V_ hotspots, potentially enhancing their packing against the juxtaposed CDR3 hotspot in amyloid (Fig. 1B, Fig. 9C). In summary, we propose that the unusual behavior of AL2 LC stems from increased exposure/dynamics observed in βC’_V_ and βE_V_ hotspots (which initiates aggregation) combined with decreased dynamics near the internal disulfide (which slows amyloid formation). Slightly enhanced amyloidogenic sequence propensity of several hotspots in AL2 *vs*. GL2 LC (Fig. 1) may also contribute to amyloid formation.

### CDR1-CDR3 link influences amyloid formation by various LCs

In all LCs explored in the current study (except for AL1_CDR1_) residue segment 25-34 overlapping CDR1 was the only segment whose protection correlated directly with overall protein stability (Figs. 2, 3, 5-8). Similarly, in prior HDX MS studies of λ6 LCs [25], local protection in CDR1 correlated directly with the protein stability; moreover, conservative substitutions in CDR1 influenced both overall protein stability and amyloid formation. A similar effect was observed in AL1_CDR1_ LC from the current study (Fig. 4). Collectively, our studies of λ6 and κ1 LCs identify CDR1 as a sensitive region that influences LC stability and amyloid formation.

The current study suggests that CDR3, which harbors four mutations increasing its amyloidogenic propensity in AL1 and in AL2, modulates amyloid formation by full-length LCs. Notably, in the stand-alone AL1 V_L_, the misfolding may be initiated by CDR3, which is more exposed/dynamic and more amyloidogenic *vs*. GL1 V_L_ (Figs. 1, 4). Interestingly, AL mutations in CDR3 were either non-destabilizing (in AL1 LC) or even stabilizing (in AL2 LC); the latter helped rescue the protein from rapid degradation (AL2 *vs*. AL2_CDR3_ LC, Figs. 7, 8). These results exemplify a general trend that moderate structural destabilization of a globular protein generally augments amyloid formation, while large destabilization counteracts amyloidogenesis and leads to rapid protein degradation [19, 32].

Our analysis of engineered LCs with amino acid substitutions in either CDR1 or CDR3 (Figs. 4-8) reveals non-additive, context-dependent effects of these substitutions, indicating interactions between CDR1 and CDR3. We posit that such interactions, facilitated by the close spatial proximity of these CDRs, can contribute to normal protein function in antigen binding and also to misfolding and aggregation in amyloid. Since CDR1 and CDR3 are disulfide-linked, the effects of AL substitutions in or near these loops on the misfolding pathway must be coupled at all steps of this pathway. Since substitutions in CDR3 and CDR1 constitute the majority of all AL mutations, we propose that mutational effects on the conformation and dynamics in CDR1 and CDR3 in native and misfolded states are among the major determinants of amyloid formation by various LCs.

### Study limitations and future directions

In the current study, the mutational effects have been interpreted by using the available atomic structures of native and fibrillary LCs. However, early misfolding intermediates – whose atomic structures are not available – are expected to be particularly sensitive to mutations and thus, likely define the metabolic fate of LCs and other amyloidogenic proteins [33]. Future studies will determine the effects of AL mutations on such intermediates. Another limitation is LC self-association, including covalent dimerization and non-covalent oligomerization, which mimics the *in vivo* effects but complicates the biophysical data interpretation. Furthermore, AL mutations may act in synergy with factors that modulate amyloid deposition *in vivo* [1], including interactions with amyloid modifiers such as glycosaminoglycans [34], LC proteolysis, or post-translational modifications [15]. Exploring the interplay of these factors with AL mutations will help bridge the gap between *in vitro* and *in vivo* mechanisms of protein misfolding.

## Supporting information

Supplementary Datafile

## Abbreviations

AL –: amyloid light chain;
CD –: circular dichroism;
CDR –: complementarity-determining region;
C_L_ –: constant domain;
HDX –: hydrogen-deuterium exchange;
GL –: germline;
LC –: immunoglobulin light chain;
MM –: multiple myeloma;
MS –: mass spectrometry;
EM –: electron microscopy;
SDS PAGE –: sodium dodecyl sulfate polyacrylamide gel electrophoresis;
ThT –: thioflavin T;
V_L_ –: variable domain.

## Acknowledgements

We thank Mari Nakamura for help with data collection using electron microscopy and Thomas Wales for technical assistance with HDX MS.

## Data availability statement

The HDX MS data have been deposited to the ProteomeXchange Consortium via the PRIDE [31] partner repository with the dataset identifier PXD039682. The summary of HDX MS experimental parameters, proteolytic maps, details of replicates, and the numeric values used to create all HDX MS figures are provided in the Supplemental Datafile 1.

