## Supplementary Appendix for "Role of Complementarity-Determining Regions 1 and 3 in Pathologic Amyloid Formation by Human Immunoglobulin κ1 Light Chains"

#### Materials and Methods

***Proteins***. The total of 14 proteins listed in Results were generated following established protocols [1]. $V_L$ domain of AL1 LC differs from GL1 in eight positions (D28G, Y32S, S63T, T74S, D92A, N93S, L94V, and P95F), $V_L$ of MM1 LC differs from GL1 in 7 positions (S30I, L33V, L46V, I48V, N53K, D92E, and F96L), and $V_L$ domain of AL2 LC differs from GL2 in 14 positions (T20I, D28T, S30R, L46V, A50G, S53N, Q55H, S56N, R61N, A84G, S91C, Y92H, T94I, and I96F). Many of these positions are in the hypervariable CDRs in or near the antigen-binding loops (Figure 1).

***SDS-PAGE.*** The protein purity and percent of covalent dimers was determined by SDS-PAGE analysis using Novex$^{TM}$ WedgeWell$^{TM}$ 10-20% Tris-Glycine Precast Gels (Thermo Fisher Scientific). Samples (~7 μg of protein) were loaded under either reducing or non-reducing conditions. Gels were stained with Coomassie Blue. Precision Plus Protein Dual Color Standards (Bio-Rad) were used as protein standards. Densitometric analysis to assess the purity and percentage of covalent dimeric species was performed using the free GelAnalyzer software (v19.1).

***Intact mass analyses.*** Intact protein mass analyses were performed using a Xevo G2 Q-TOF mass spectrometer (Waters) and a nanoAcquity UPLC system (Waters). Each protein sample (~70 pmols, 2 μL, in 10 mM sodium phosphate, 150 mM NaCl, pH 7.5) was treated with 100 mM TCEP for 20 min at 40 °C, followed by the addition of 45 μL of 0.1% formic acid immediately before injection. Separations were carried out using an in-house packed POROS 20-R2 trap eluted with a 15%–70% gradient of acetonitrile at 100 μL/min over 6 minutes. Mass measurements were obtained in positive ion mode, with the capillary potential set to 3.2 kV and source temperature to 80 °C. Average mass values were determined using MagTran 1.02 software [2]. The instrument was regularly calibrated with horse heart myoglobin such that the maximum error of mass determination was +/- 1.0 Da.

***Limited proteolysis.*** Samples containing 0.25 mg/mL protein were digested at 37 °C with trypsin at an enzyme:substrate weight ratio of 1:100. The reaction was quenched at specified time points using 1 mM phenylmethylsulfonyl fluoride. The products were analyzed by SDS PAGE using 12% Tris - Glycine gel stained with Coomassie Blue.

***Circular dichroism (CD).*** Far-UV CD spectra and the melting data were recorded using a Jasco J-1500 spectropolarimeter (Jasco Inc., Japan). CD spectra were recorded at 195-250 nm using samples containing 0.3 mg/mL protein in 1 mm quartz cells. After the baseline subtraction, the spectra were smoothed using a noise reduction routine. The data were normalized to protein concentration and expressed in units of molar residue ellipticity (MRE). Melting data were recorded at 206 nm to monitor protein unfolding during heating from 25 °C to 65 °C at a constant rate of 0.5 °C/min. The melting data were normalized to fraction unfolded and approximated by a sigmoidal function.

***Fluorescence spectroscopy.*** Formation of amyloid-like structure was monitored in real time by thioflavin T (ThT) binding and fluorescence using TECAN microplate reader (Life Sciences, Switzerland). The samples containing 0.3 mg/mL protein and 10 μM ThT were subjected to double-orbital shaking at 37 °C. ThT emission ($\lambda_{ex}$ = 450 nm, $\lambda_{em}$ = 482 nm) was recorded at 10 min intervals for up to 140 h. The emission of ThT in buffer alone was subtracted.

All experiments were performed in technical triplicates of biological triplicates.

***Transmission electron microscopy (EM).*** For electron microscopy, sample at ~0.5 mg/ml in PBS was diluted 4x in 50 mM Tris pH 7.6. A 5 µl drop was deposited onto a formwar/carbon-coated mesh-300 copper grid for 3 min. The grids were washed on three drops of 50 mM Tris pH 7.6, stained on two drops of 1% uranyl acetate and blotted. Images were recorded at a magnification of 25,000 or 45,000 x using a CM12 transmission electron microscope (Philips Electron Optics, the Netherlands) operated at 100 kV and equipped with a Tietz 2Kx2K CCD camera (TVIPS, Germany).

***Hydrogen deuterium exchange mass spectrometry (HDX MS).*** The recommended summary [3] of HDX MS experimental parameters, proteolytic maps for all proteins, details of replicates acquired, and the numeric values used to create all HDX MS figures are provided in the Supplemental Datafile 1. The HDX MS data have been deposited to the ProteomeXchange Consortium via the PRIDE [4] partner repository with the dataset identifier PXD039682.

Deuterium labeling: $\kappa$1 IgG LCs aliquots at 30 µM (stock concentration) were freshly prepared in biological replicates prior to HDX MS analysis in 10 mM sodium phosphate, 150 mM NaCl, pH 7.5, $H_2O$. The starting stock solution of protein was 30 µM, of which 1 µL was mixed with 18 µL of labeling buffer (10 mM sodium phosphate, 150 mM NaCl, pD 7.5, 99.9% $D_2O$) to initiate labeling at 20 °C. After each labeling time (10 seconds, 1 minute, 10 minutes, 1 hour, 4 hours), the labeling reaction was quenched with the addition of 30 µL of ice-cold quenching buffer (7 M Gdn·HCl, 0.6 M TCEP, 0.8% formic acid, pH 2.4, $H_2O$). Each quenched solution was kept in an ice bath for 30 seconds before being diluted with 50 µL of 0.8% formic acid (final volume 99 µL) and then injected into the LC MS system. Fully deuterated samples for back-exchange correction were prepared as described in [5]. Briefly, protein solutions (15 µL, 30 µM) were speed vac to dryness, resuspended in 7 M Gdn·HCl containing 50 mM DTT (15 µL), and heated at 90 °C for 5 min. After cooling to 20 °C, 18 µL of labeling buffer were added to 1 µL of denatured protein solution, and the exchange reaction was allowed to proceed at 50 °C for 10 min. The sample was cooled to 0 °C, quenched with the addition of 30 µL of ice-cold quenching buffer, kept at 0 °C for 30 seconds, diluted with 50 µL of 0.8% formic acid and analyzed immediately by LC MS.

LC / MS: All LC steps were performed with a Waters HDX system containing an HDX unit and two Acquity I-class UPLC pumps. Deuterated and control samples were digested online in the HDX cooling unit, where the digestion chamber was held at 15 °C, using an Affipro Nepenthesin-2 column (2.1 mm × 20 mm). Peptides were trapped and desalted on a VanGuard Pre-Column trap [2.1 mm × 5 mm, ACQUITY UPLC BEH C18, 1.7 µm (Waters, 186002346)] for 3 minutes at 100 µL/min. Peptides were eluted from the trap using a 5%–35% gradient of acetonitrile over 6 minutes at a flow rate of 100 µL/min using an ACQUITY UPLC HSS T3, 1.8 µm, 1.0 mm × 50 mm column (Waters, 186003535). The main cooling chamber of the Waters HDX unit, which housed all the chromatographic elements, was held at 0.0 ± 0.1 °C for the entire time of the measurements. The back pressure averaged ~9,000 psi at 0 °C and 5% acetonitrile 95% water. To eliminate peptide carryover, a wash solution (1.5 M guanidinium chloride, 0.8% formic acid and 4% acetonitrile) was injected into the Nepenthesin-2 column during each analytical run. Mass spectra were acquired using a Waters Synapt XS HDMS$^E$ mass spectrometer in ion mobility mode. The mass spectrometer was calibrated with direct infusion of a solution of glu-fibrinopeptide (Sigma, F3261) at 200 femtomole/µL at a flow rate of 5 µL/min prior to data collection. A conventional electrospray source was used, and the instrument was scanned over the range 50 to 2000 m/z. The instrument configuration was: capillary voltage 2.5 kV, trap collision energy at 4 V, sampling cone at 35 V, source temperature of 80 °C, and desolvation temperature of 175 °C. The error of determining the deuterium levels was ± 0.20 Da in this experimental setup.

Data processing: Peptides were identified from replicate HDMS$^E$ analyses (see Supplemental Datafile 1) of undeuterated control samples using PLGS 3.0.1 (Waters, 720001408EN). Peptide masses were identified from searches using non-specific cleavage of a custom database containing the sequence of κ1 GL1 $V_L$ (IGKVLD-33*01), GL2 $V_L$ (IGKVLD-39*01), and $C_L$ (IGKCLD) (UniProt: P01593, P01597, and P01834, respectively), no missed cleavages, no PTMs, a low energy threshold of 135, an elevated energy threshold of 20 and an intensity threshold of 500. No false discovery rate (FDR) control was performed. The peptides identified in PLGS (excluding all neutral loss and in-source fragmentation identifications) were then filtered in DynamX 3.0.1 (Waters, 720005145EN) implementing a minimum products per amino acid cut-off of 0.25, at least 2 consecutive product ions. Those peptides meeting the filtering criteria were further processed automatically by DynamX followed by manual inspection of all spectra and data processing steps. The relative amount of deuterium in each peptide was determined by subtracting the centroid mass of the undeuterated form of each peptide from the deuterated form, at each time point, for each condition. Correction for back exchange to report percent deuteration (%D) was done as described [5] according to the equation $\%D = [(m_t – m_0) / (m_{maxD} – m_0)]$, where $m_t$ is the observed peptide centroid mass at a given labeling time point t, $m_0$ is the undeuterated peptide centroid mass, and $m_{maxD}$ is the maximally deuterated peptide centroid mass. Percent deuteration values were used to generate %D uptake graphs and skyline plots. Accordingly, all data plotted on tertiary structures was likewise back exchange corrected. All measured and processed values can be found in Supplemental Datafile 1.

***Amyloid protein analysis by SDS PAGE and mass spectrometry.*** To determine the protein composition in amyloid fibrils formed *in vitro*, AL1 LC was incubated with trypsin for 5 days under amyloid-promoting conditions (37 °C with shaking) and the aggregated material was spun down and analyzed. Non-reducing SDS PAGE showed the major band at ~10 kDa which was excised and analyzed for protein composition. Gel slice was rinsed with a 1:1 acetonitrile : water on an orbital shaker for 15 min at 22 °C, dehydrated in acetonitrile, and dried in a vacuum centrifuge. Next, 10 mM dithiothreitol (DTT) in 100 mM $NH_4HCO_3$ was added, the proteins were reduced for 1 h at 56 °C, cooled to 22 °C, and the solution was replaced with 55 mM iodoacetamide in 100 mM $NH_4HCO_3$. After 45 min incubation at 22 °C in the dark with occasional vortexing, the gel slice was washed with 100 mM $NH_4HCO_3$ for 10 min, dehydrated by acetonitrile, rehydrated in 100 mM $NH_4HCO_3$, and dehydrated again. The liquid phase was removed and the gel piece was dried.

For enzymatic cleavage, 100 μl of 50 mM $NH_4HCO_3$ and 5 μl of endoproteinase Lys-C (NEB 0.1 μg/μl) was used in an ice-cold bath. After 45 min, 50 μl of 50 mM $NH_4HCO_3$ was added, and the samples were incubated overnight at 37 °C. Peptides were extracted by one change of 20 mM $NH_4HCO_3$ and one change of 0.5% trifluoro-acetic acid (TFA) in 50% acetonitrile at 22 °C.

For matrix-assisted laser desorption/ionization (MALDI) analysis, the dried peptides were reconstituted in 20 μl of 0.1% TFA, and 10 μl taken for desalting using Millipore UC18 zip-tips. Peptides were eluted from the tip using Sigma alpha-cyano-4-hyrdoxycinnamic acid (10 mg/ml in 0.1% TFA). MALDI analysis was performed using an AB-Sciex 4800 MALDI-TOF/TOF instrument in positive reflection mode and MSMS of selected peptides were obtained using a 2 kV pos mode method. The peptide identity and coverage were determined using Expasy Pepfind software.

| Protein | Intact MS analyses Theor. | Exper. | Δ | SDS PAGE analyses Purity (%) | Covalent Dimer (%) |
|---|---|---|---|---|---|
| AL1_LC | 24335.1 | 24334.0 | -1.1 | 93.5 | 11.7 |
| AL1_VL | 12744.2 | 12743.1 | -1.1 | 98.4 | N/A |
| GL1_LC | 24504.2 | 24503.0 | -1.2 | 98.8 | 8.6 |
| GL1_VL | 12913.3 | 12912.2 | -1.1 | 99.9 | N/A |
| MM1_LC | 24482.3 | 24481.0 | -1.3 | 95.0 | 29.0 |
| CR | 12819.2 | 12818.2 | -1.0 | 99.9 | 54.5 |
| AL1_CDR1 | 24469.2 | 24467.8 | -1.4 | 91.3 | 13.6 |
| AL1_CDR3 | 24370.1 | 24368.7 | -1.4 | 98.8 | 10.0 |
| GL1_CDR3 | 24469.2 | 24467.9 | -1.3 | 90.0 | 25.7 |
| AL2_LC | 24426.3 | 24425.8 | -0.5 | 95.7 | 18.6 |
| AL2_CDR3 | 24390.2 | 24389.0 | -1.2 | 85.1 | 8.0 |
| GL2_LC | 24350.1 | 24349.8 | -0.3 | 99.9 | 20.4 |
| GL2_CDR3 | 24386.2 | 24384.8 | -1.4 | 96.4 | 14.8 |

**Table S1.** Characterization of all κ1 LCs investigated by Intact MS and SDS PAGE analysis. Intact MS analysis was performed under reducing conditions (see Methods in Supplementary Appendix). Theoretical (Theor.) and measured experimental (Exper.) average mass values and their differences (Δ) are reported in Daltons for each protein form. SDS PAGE analysis was performed both under reducing conditions, to assess the purity of each protein batch, and under non-reducing conditions, to estimate the percentage of the covalent dimeric form present in solution. The values (in percentage) were derived from gels in Fig. S1 by using Gel analyzer software. Further details can be found in the Supplementary Appendix.

#### Supplementary Figures

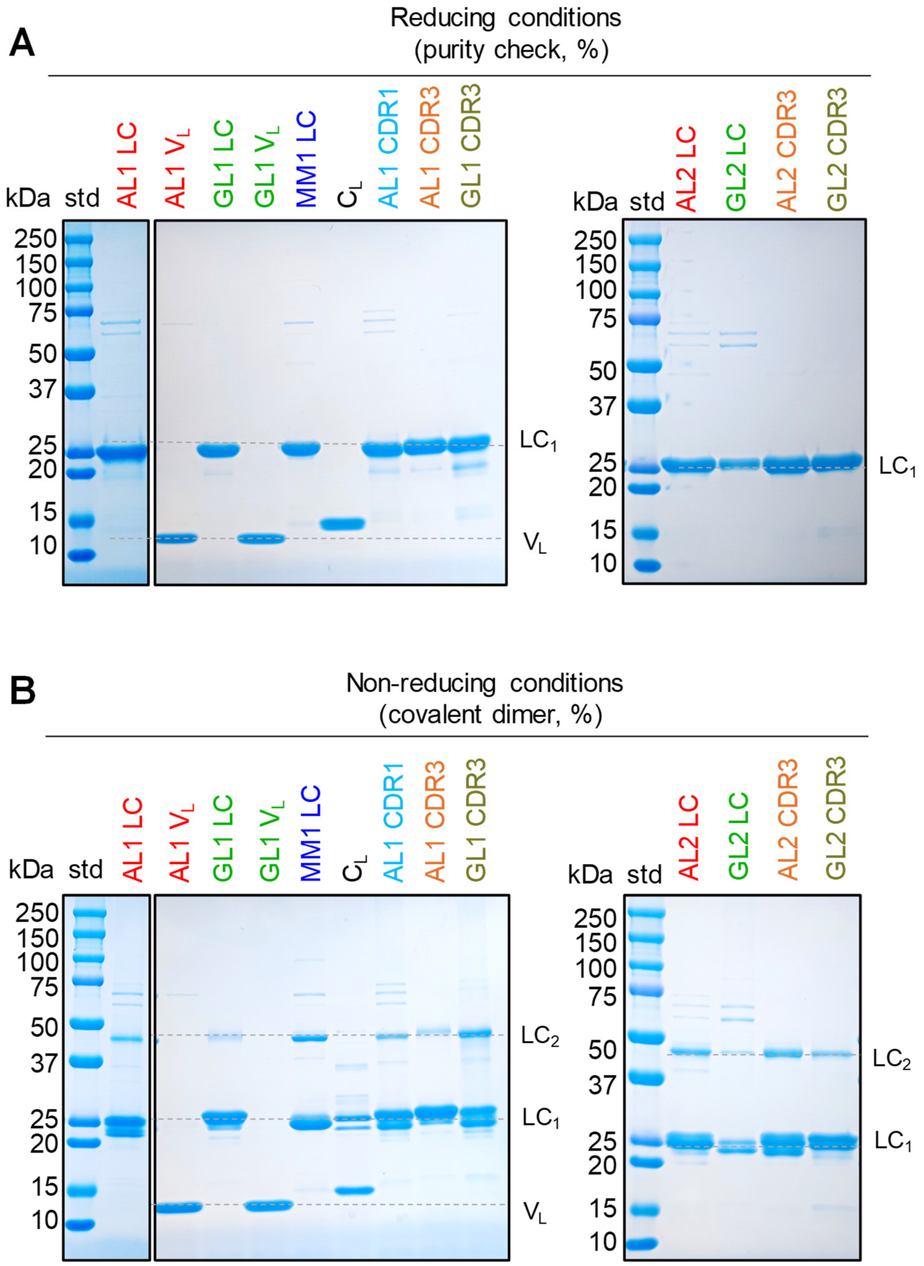


**Figure S1**. Chemical and conformational characterization of κ1 LC proteins. SDS PAGE analysis under reducing (**A**) and non-reducing conditions (**B**) of LCs and their isolated $V_L$ and $C_L$ domains. Gels were run by loading 7 mg of protein per well. Gels bands corresponding to the covalent LC homodimer ($LC_2$), the monomer ($LC_1$), and the stand-alone $V_L$ are indicated. The purity and percentage of covalent dimer calculated from these gels are listed in Table S1.

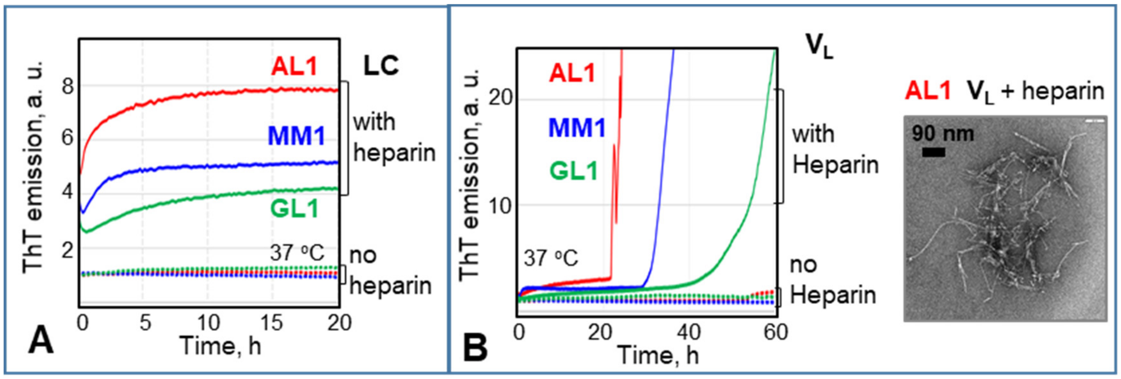


**Figure S2.** Amyloid formation in the presence of heparin by the patient-derived LCs related to germline gene *IGKVLD-33*01*. The proteins were incubated at 37 °C with shaking as described in Methods. Formation of amyloid-like structure was monitored continuously by ThT emission at 482 nm. Representative data for each protein are color-coded: AL1 (red), MM1 (blue), GL1 (green). The time course of amyloid formation is shown for full-length LCs (**A**) and their stand-alone $V_L$ domains (**B**), along with a representative transmission electron micrograph (negative stain) of AL1 $V_L$ after 50 h of incubation.

**Figure S3** (next page)**.** Skyline plots at all HDX MS time points for patient-derived full-length LCs and their stand-alone domains. Full-length LCs of AL1, GL1, and MM1 (solid symbols), stand-alone $V_L$ domains of AL1 and GL1 (open, colored symbols), and their common stand-alone $C_L$ domains (open black symbols). The data are shown for the peptide fragments common to all proteins. The exchange times are indicated. Linear representation of the native secondary structure in κ1 LC and predicted amyloid hotspots that have the AmylPred2 consensus number ≥5 (purple) are shown at the top.

### Figure S3

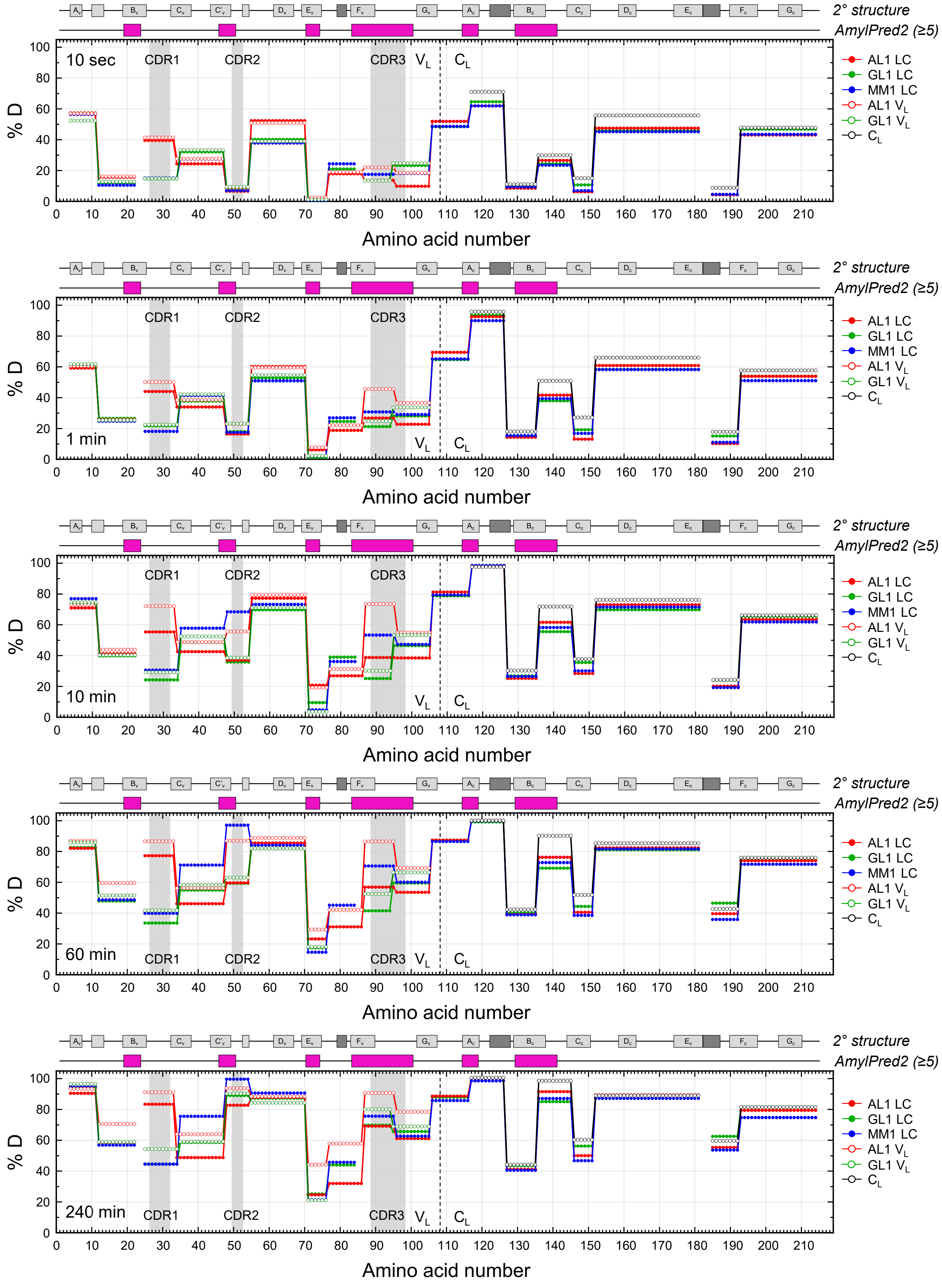
2° structure
AmylPred2 (≥5)
% D
Amino acid number
10 sec
1 min
10 min
60 min
240 min
CDR1
CDR2
CDR3
$V_L$
$C_L$
AL1 LC
GL1 LC
MM1 LC
AL1 $V_L$
GL1 $V_L$
$C_L$

**Figure S4** (next page). Skyline plots at all HDX MS time points for engineered full-length LCs. Data for AL1$_{CDR1}$ (teal), AL1$_{CDR3}$ (orange) and GL1$_{CDR3}$ (olive) are compared with those of AL1 (red) and GL1 (green) LCs. The exchange times are indicated. Linear representation of the native secondary structure in $\kappa$1 LC and predicted amyloid hotspots that have the AmylPred2 consensus number ≥5 (purple) are shown at the top.

### Figure S4

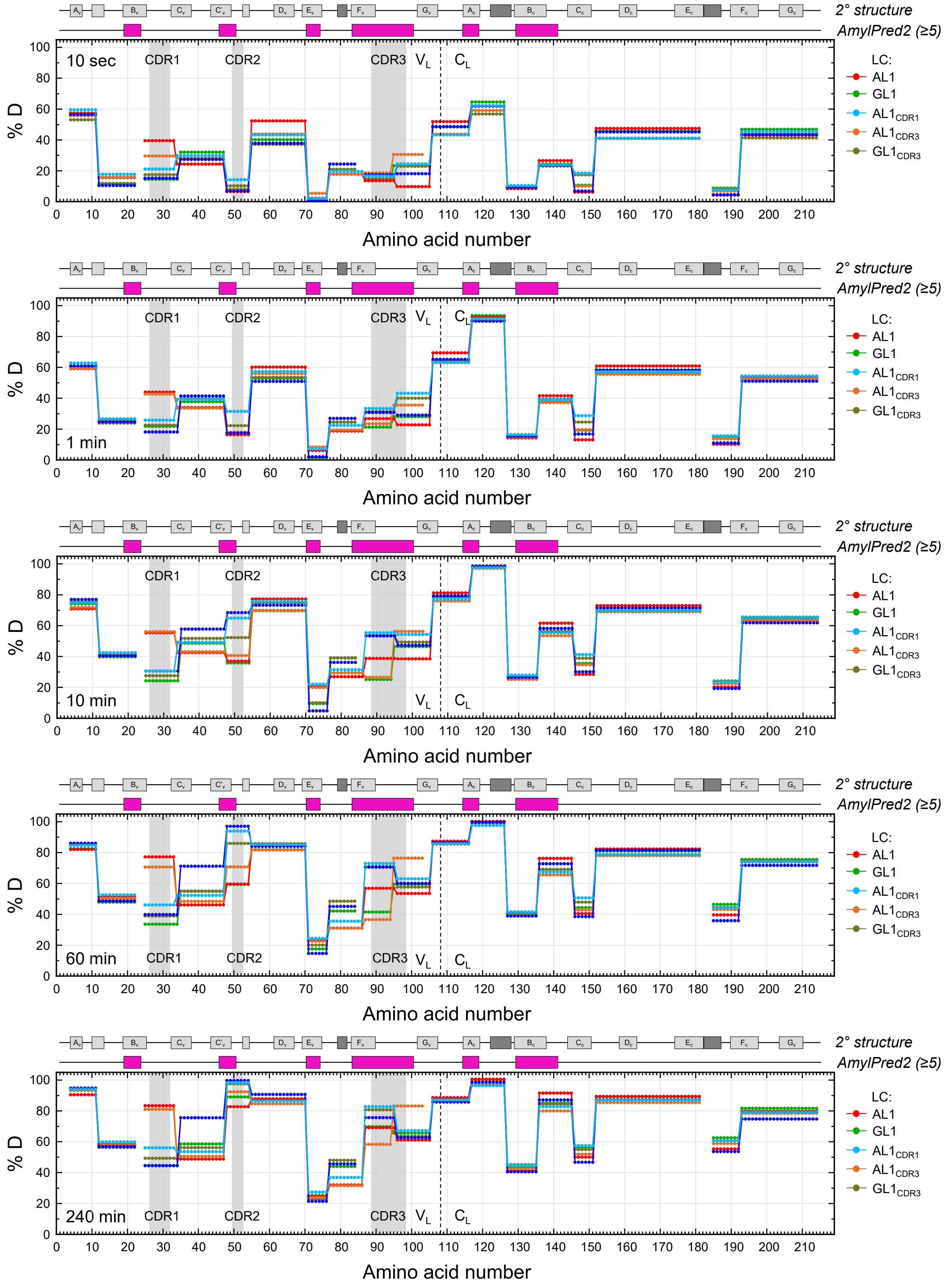
2° structure
AmylPred2 (≥5)
10 sec
1 min
10 min
60 min
240 min
CDR1
CDR2
CDR3
$V_L$
$C_L$
% D
Amino acid number
LC:
AL1
GL1
$AL1_{CDR1}$
$AL1_{CDR3}$
$GL1_{CDR3}$

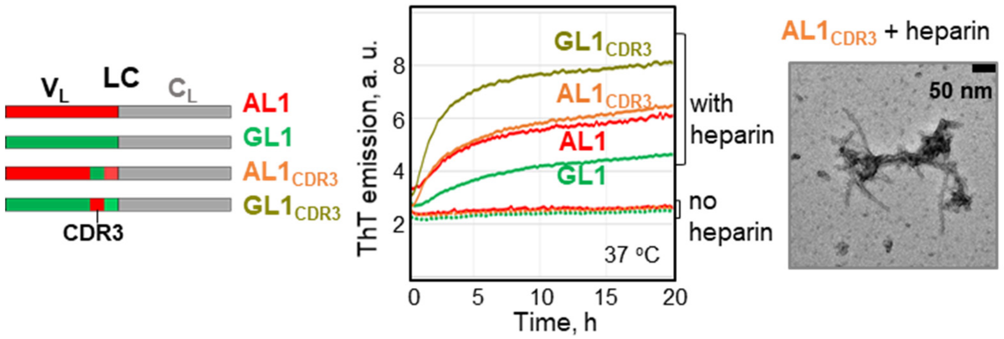


**Figure S5.** Amyloid formation in the presence of heparin by full-length LCs related to germline gene *IGKVLD-33*01*: comparison of engineered proteins, $GL1_{CDR3}$ (olive) and $AL1_{CDR3}$ (orange), with GL1 LC (green) and AL1 LC (red). Schematics on the left illustrates the primary structures of the engineered proteins. The time course of amyloid formation monitored by ThT fluorescence is shown along with a representative transmission electron micrograph of AL1 $LC_{CDR3}$ after 50 h of incubation. Experimental conditions are described in Methods.

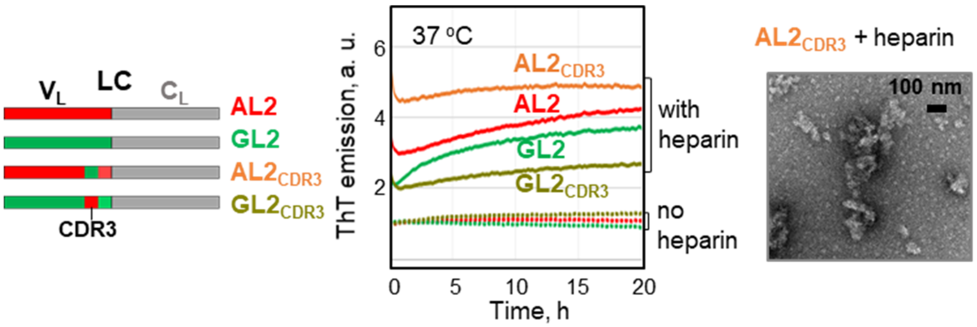


**Figure S6**. Amyloid formation in the presence of heparin by the patient-derived and engineered full-length LCs related to the germline gene *IGKVLD-39*01*: AL2 (red), GL2 (green), $GL2_{CDR3}$ (olive) and $AL2_{CDR3}$ (orange). Schematics on the left illustrates the primary structures of the engineered proteins. The time course of amyloid formation monitored by ThT fluorescence is shown together with an electron micrograph (negative stain) of the sample after 50 h incubation under amyloid-promoting conditions. Experimental conditions are described in Methods.

**Figure S7** (next page). HDX MS skyline plots for all time points of part 2 LCs (AL2, GL2, $AL2_{CDR3}$, $GL2_{CDR3}$). Linear representation of the native secondary structure in κ1 LC and predicted amyloid hotspots that have the AmylPred2 consensus number ≥5 (purple) are shown at the top.

#### Figure S7

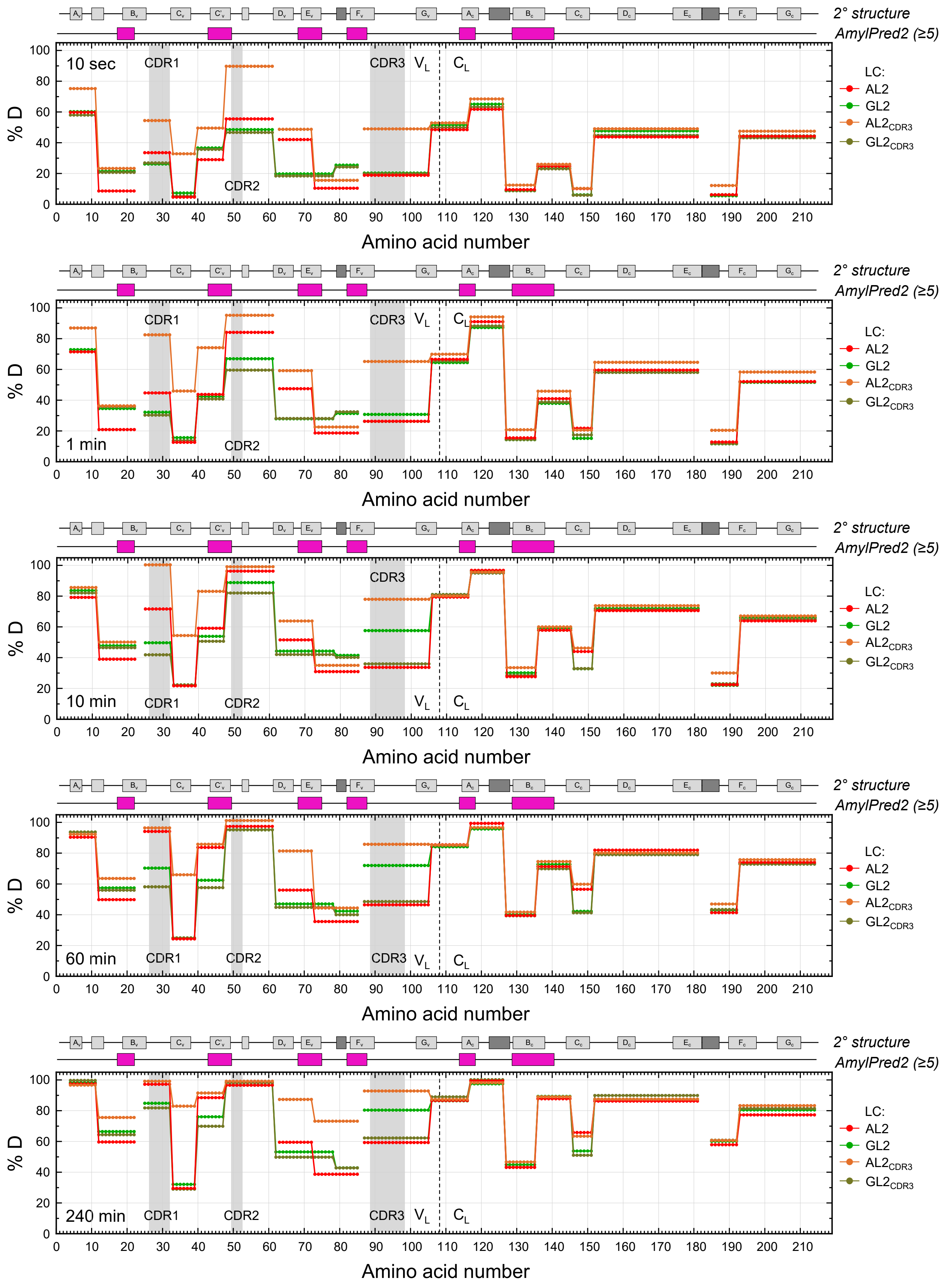
2° structure
AmylPred2 (≥5)
10 sec
1 min
10 min
60 min
240 min
CDR1
CDR2
CDR3
$V_L$
$C_L$
% D
Amino acid number
LC:
AL2
GL2
$AL2_{CDR3}$
$GL2_{CDR3}$

**Figure S8** (following pages). Additional mapping of HDX MS data onto tertiary structures at all time points, intended to supplement Fig 3C-D, Fig. 6C, and Fig. 8C.

### Figure S8

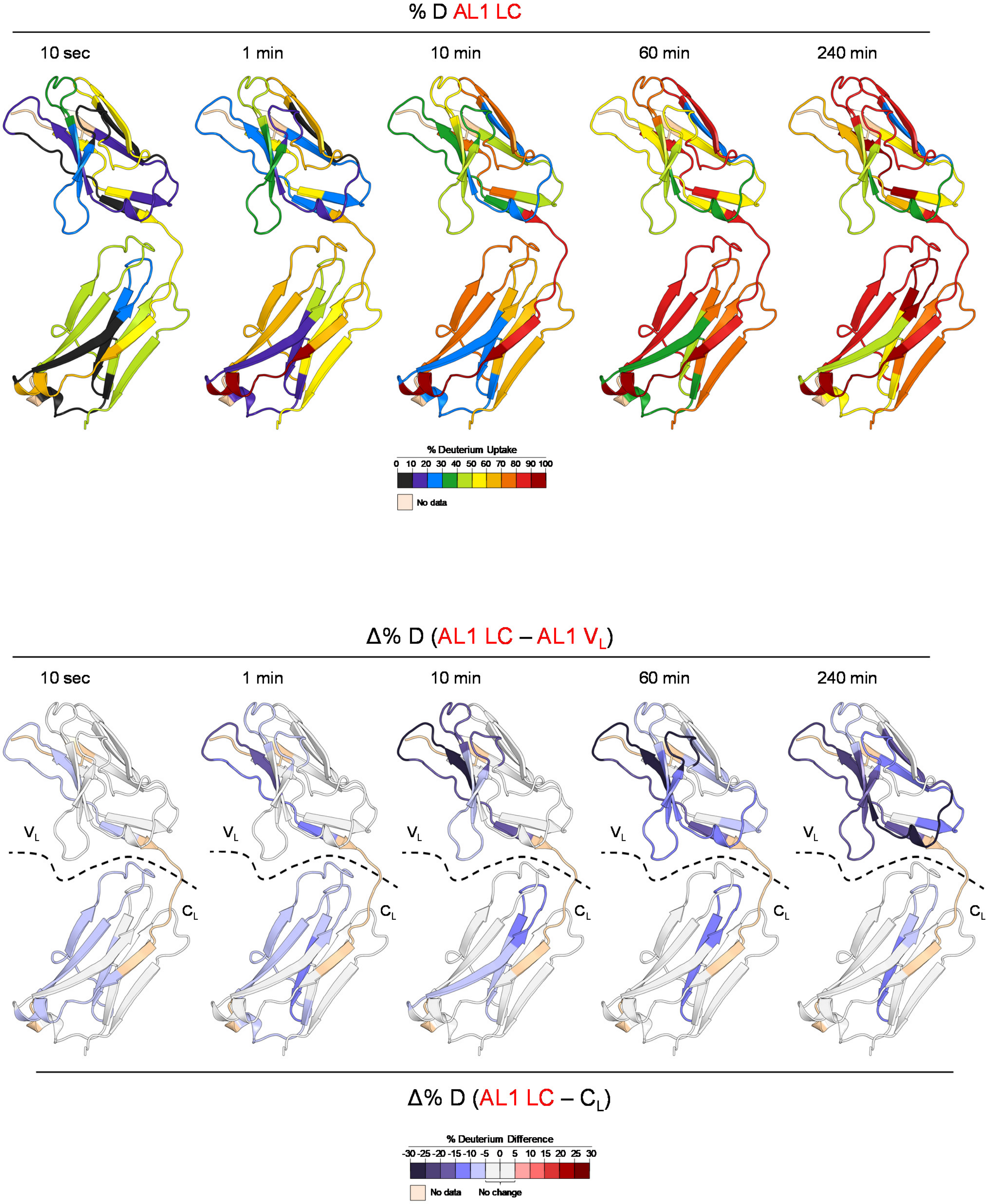
% D AL1 LC
10 sec
1 min
10 min
60 min
240 min
% Deuterium Uptake
0 10 20 30 40 50 60 70 80 90 100
No data
Δ% D (AL1 LC – AL1 V_L)
10 sec
1 min
10 min
60 min
240 min
V_L
C_L
Δ% D (AL1 LC – C_L)
% Deuterium Difference
-30 -25 -20 -15 -10 -5 0 5 10 15 20 25 30
No data
No change

### Figure S8, continued

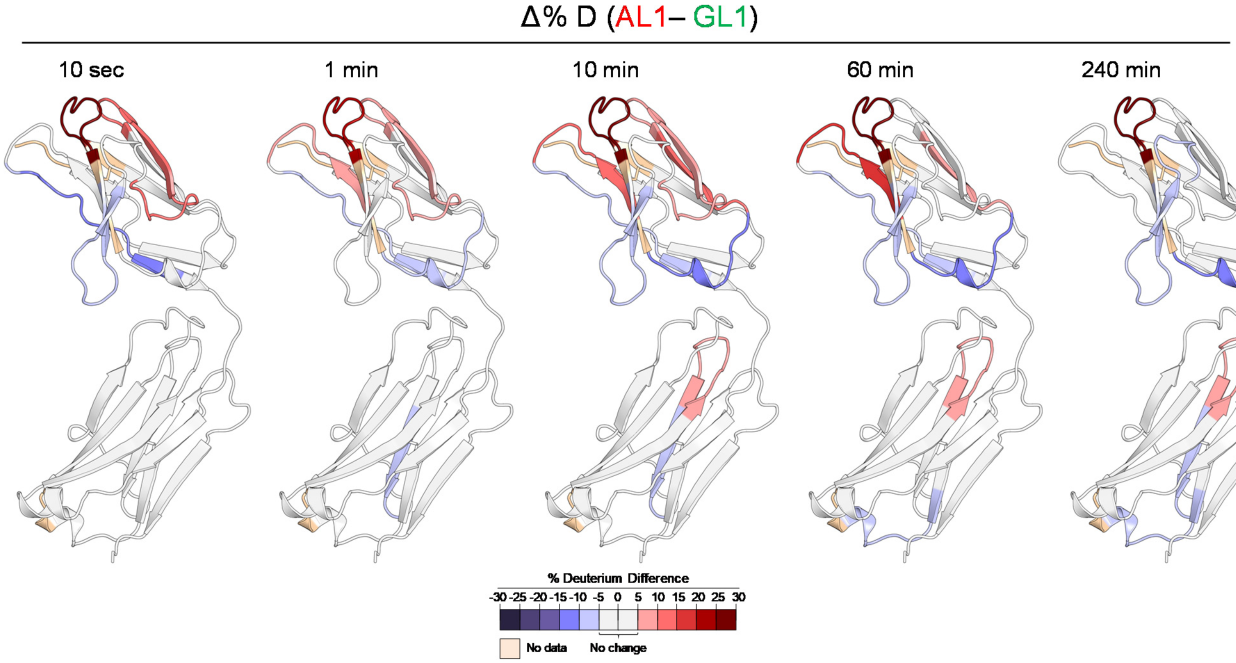


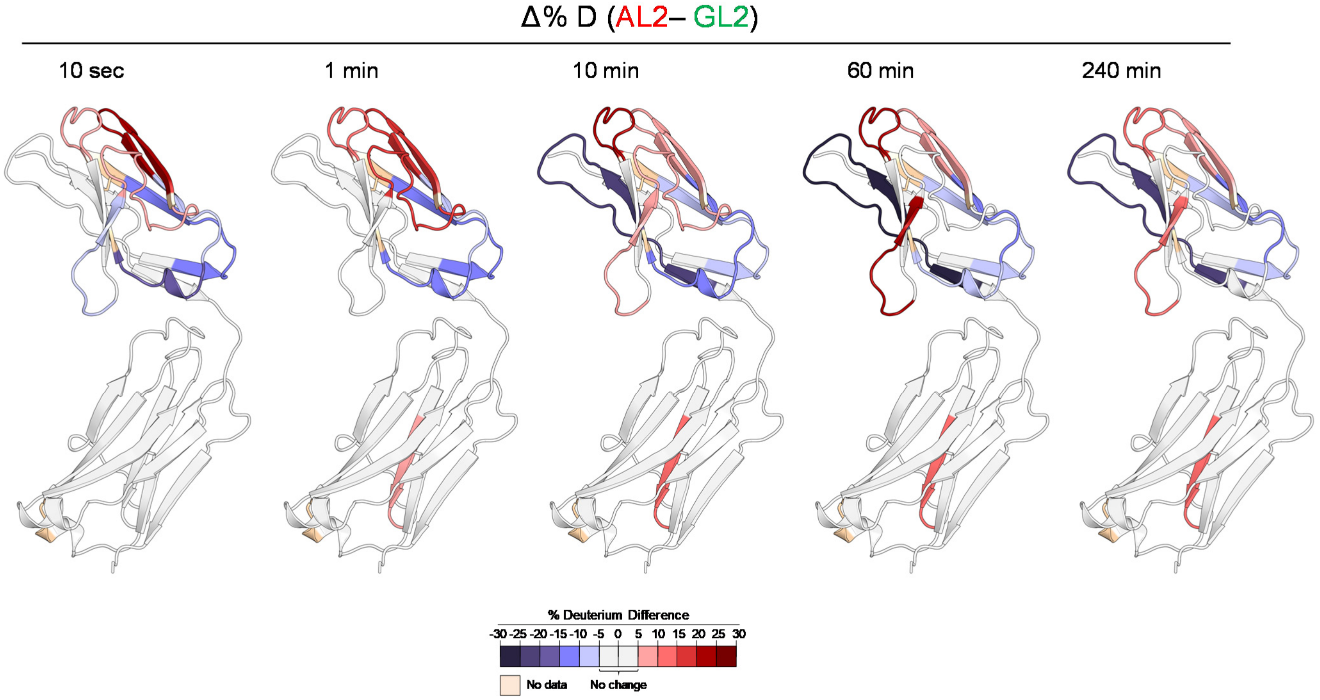
